# Evaluating the legacy of drought exposure on root and rhizosphere bacterial microbiomes over two plant generations

**DOI:** 10.1101/2025.09.19.677321

**Authors:** A. Fina Bintarti, Abby Sulesky-Grieb, Joanna Colovas, Brice Marolleau, Tristan Boureau, Marie Simonin, Matthieu Barret, Ashley Shade

## Abstract

Drought is a critical risk in developing countries for staple crops like common bean (*Phaseolus vulgaris* L.). We conducted an experiment to understand the legacy effects of repeated drought exposure across plant generations on the root and rhizosphere microbiome of the common bean, hypothesizing that a legacy of exposure improves overall plant microbiome resilience. We profiled the bacterial microbiome using marker gene amplicon sequencing over two plant generations in a complete factorial design for two common bean genotypes, Red Hawk and Flavert. We performed parallel experiments for Red Hawk in two different countries using soils of Pays de la Loire, France, and Michigan, USA. Despite the clear and relatively consistent drought effects on the plant phenotypes, there was neither response of the Red Hawk microbiomes to drought, nor a notable legacy of drought exposure. For Flavert, there was a minor legacy drought effect for the second generation in the rhizosphere microbiome beta diversity. This study demonstrates that rhizosphere microbiomes can be resistant to drought stress and that cross-generational legacy depends on soil origin and host genotype. Such parallel experiments across countries, while difficult to implement, are useful to inform generalities and build theory towards prediction on microbiome responses to global change.

## Introduction

Contending with severe and sometimes unpredictable environmental stress is now standard for agricultural systems (Furtak and Wolińska 2023; Maity *et al*. 2023; Semeraro *et al*. 2023). Changing weather patterns have resulted in droughts in many regions, which are particularly devastating in developing countries that face food insecurity (Gourdji *et al*. 2015; Madadgar *et al*. 2017; Ackerl *et al*. 2023; Chivangulula, Amraoui and Pereira 2023; Ouko and Odiwuor 2023). An important and growing area of research is focused on the impact of these environmental changes on crops, with particular interest in the plant microbiome (Shade 2018; Cheng, Zhang and He 2019; Hartman and Tringe 2019; Trivedi *et al*. 2022). The plant microbiome comprises millions of bacterial, archaeal, and fungal cells that inhabit the plant’s surfaces, tissues, and root structures (Compant *et al*. 2019; Trivedi *et al*. 2020). The beneficial members of the microbiome play an essential role in plant health in a changing climate (Goh *et al*. 2013; Lata *et al*. 2018; Bokhari *et al*. 2019). They collectively also provide vital agricultural services, including water and nutrient assimilation to the plant, nutrient cycling in the soil and root system, and pathogen resistance (Compant *et al*. 2019; Trivedi *et al*. 2020; Noman *et al*. 2021; Santos and Olivares 2021).

Many studies have improved our knowledge of the plant microbiome and its role for various crops (Walters *et al*. 2018; Chen *et al*. 2019; Liu *et al*. 2019). While some studies have addressed the impacts of environmental stress on the microbiome of plants (Santos-Medellín *et al*. 2017; Mavrodi *et al*. 2018; Timm *et al*. 2018; Veach *et al*. 2020; Wang *et al*. 2020; Russell and McFrederick 2022; Tiziani *et al*. 2022), few have studied this over multiple generations of repeated stress exposure (Rodríguez *et al*. 2023; Sulesky-Grieb *et al*. 2024). With the continuing trends of climate change, repeated exposure to drought could become standard for many agricultural areas that have historically faced droughts less frequently. These repeated stress conditions over multiple growing seasons and plant generations may create unique challenges for crops, such as negative impacts on seed quality and long-term alterations of soil conditions (Furtak and Wolińska 2023; Maity *et al*. 2023).

Our study aimed to understand the implications of repeated drought on the microbiome of the legume common bean (*Phaseolus vulgaris* L.). Legumes provide valuable nutrition and perform symbiotic nitrogen fixation, providing usable nitrogen to other crops (Stacey 2007). Legume services are particularly valuable in developing economies that are most significantly impacted by environmental stress due to a lack of infrastructure to manage poor growing conditions and reliance on subsistence farming. Common beans are a vital food source worldwide, with 27.5 million metric tons of food produced annually and cultivated across 34.8 million hectares of land as of 2020 (Castro-Guerrero *et al*. 2016; Uebersax *et al*. 2023). *Phaseolus vulgaris* L. includes a wide range of “dry edible bean” and fresh “garden bean” crops, including kidney beans, black beans, navy beans, green beans, and others (Graham and Ranalli 1997). These varieties have been selected over 8,000 years of domestication from two ancestral lines in Mesoamerica and the Andes Mountains (Schmutz *et al*. 2014; Castro-Guerrero *et al*. 2016). Furthermore, the ability to continue to produce dry beans in developing economies is vital for food security (Shrestha and Nepal 2016).

Breeding has been done to improve the drought tolerance of common bean varieties, but managing the plant microbiome may also offer some options to improve plant performance during drought (Sofi *et al*. 2021; Petrushin, Vasilev and Markova 2023). Various studies have been conducted to understand the bean rhizosphere (Pérez-Jaramillo *et al*. 2017; Stopnisek and Shade 2021) and seed (Klaedtke *et al*. 2016; Chesneau *et al*. 2020; Bintarti *et al*. 2022b) microbiome, and its association with nitrogen-fixing rhizobia (Da Silva, Tsai and Bonetti 1993; Grange *et al*. 2007; De Ron *et al*. 2017), as well as the impact of abiotic stress, such as drought, on the bean microbiome (Bintarti *et al*. 2022a; Bandopadhyay *et al*. 2024; Sulesky-Grieb *et al*. 2024). However, it remains unclear how repeated drought across generations may affect the below ground microbiome of common bean.

To address this knowledge gap, we conducted a two-generation experiment in which we exposed common bean plants to drought in agricultural soil under controlled growth chamber conditions. We first aimed to confirm that (1) under our experimental conditions, the drought negatively impacts the bean plant health, and then hypothesized that (2) within a generation, the drought reduces the bacterial root and rhizosphere microbiome alpha diversity and increases its variance, and (3) that the repeated exposure to drought over two generations has compounded, legacy consequences for the plant and its microbiome. To assess the impact of drought on the species more broadly, we tested our hypotheses in two different genotypes of *Phaseolus vulgaris* L., the Red Hawk variety, a kidney bean cultivar bred in North America, and Flavert, a European Flageolet bean cultivar (Kelly *et al*. 1998; Bitocchi *et al*. 2017; Carrère *et al*. 2023). We included production soils from two distant geographic locations where beans are grown, Pays de la Loire, France, and Michigan, USA, to determine how the location shapes drought response.

The experiment started with seeds from a Generation 0 (G0) seed pool, which we germinated and then exposed to either a well-watered condition or a drought condition in which water was decreased by 66% (Fig. S1.1). We harvested seeds from Generation 1 (G1) plants and used them to grow Generation 2 (G2), in which the plants either received the same or opposite treatment as in G1, with the parent lines tracked and distributed across the G2 experimental conditions. The plant growth, yield, and 16S V4 rRNA gene amplicon sequencing of the root and rhizosphere microbiome were analyzed to test our hypotheses.

## Materials and Methods

### Bean cultivars

*Phaseolus vulgaris* L. var. Red Hawk, was selected as a dry bean variety (Kelly *et al*. 1998) common in Michigan production and var. Flavert was selected as a Flageolet bean variety common in French production (Carrère *et al*. 2023). Red Hawk seeds in Michigan were obtained from the Michigan State University Bean Breeding Program from their 2019 harvest and stored at 4°C until ready for use in experiments. Flavert and Red Hawk seeds in France were purchased from Vilmorin-Mikado (Limagrain group, France). These seeds were designated as G0 seeds and germinated to produce G1 plants.

### Field soil preparation

For G1 and G2 experiments performed in Michigan, agricultural field soil was collected from the Michigan State University Agronomy Farm from a field that had grown common bean in 2019 (42°42’57.4”N, 84°27’58.9”W, East Lansing, MI, USA). Field soil was used because it was expected to contain the legacy microbiome from the most recent bean crop, and thus represent more authentically the microbiome and soil conditions for bean production. The soil was a sandy loam with an average pH of 7.2 and organic matter content of 1.9%, as assessed by the Michigan State University Soil Plant and Nutrient Laboratory by their standard protocols. The soil was collected before each planting group from the same field location. The 1-3 cm dry top layer of soil and plant debris was avoided. Soil was stored covered at 4°C until the experiment. Immediately before planting, the soil was passed through a 4 mm sieve to remove rocks and plant debris, and the soil was mixed with autoclaved coarse vermiculite at a 50% v/v ratio.

For G1 experiments performed in Pays de la Loire, the soil was collected from the experimental station of the National Federation of Seed Multipliers (FNAMS, 47°28’012.42” N – 0°23’44.30” W, Brain-sur-l’Authion, France) where common bean had been cultivated in 2016. This soil was a clay sand limestone with pH 7.1 and 1.9% organic matter content. The soil was sieved and mixed with coarse vermiculite using the abovementioned method. Soil for G2 was collected from the experimental fields belonging to the Institut National de Recherche pour l’Agriculture, l’Alimentation et l’Environnement (INRAE) in Angers, France (47°28’50.7”N, 0°36’31.4”W). A different soil was used in G2 to simulate a realistic scenario during which seed multiplication occurs at a site (seed producer) different from the bean production (farm). This soil had a sandy-loam texture with a pH of 6.5 and 1.9% organic matter content. The soil was sieved and mixed with vermiculite in the same fashion as G1.

### Surface sterilization and seed germination

For each Michigan planting group, Red Hawk seeds were surface sterilized with a solution of 10 % bleach and 0.1 % Tween20 before planting. Seeds were randomly selected from the G0 seed supply for G1 and from the harvested G1 seeds for G2. We avoided visibly cracked or moldy seeds. Seeds were placed in a Petri dish lined with sterile filter paper and 1-2 mL sterile deionized (DI) H_2_O was applied to the filter paper. Petri dishes were stored in the dark at room temperature for 3-4 days for seeds to germinate, with an additional 2 mL sterile DI H_2_O added halfway through the germination period. Once seeds had sprouted radicle roots, they were transferred to the pots. Seeds planted in Pays de la Loire were germinated directly in the prepared pots of soil without prior surface sterilization. The Pays de la Loire experiment included both Red Hawk and Flavert cultivars under the same soil and treatment conditions for each generation. After G1, seeds were removed from the pods of G1 plants, pooled by each plant in 50 mL conical tubes, and stored at 4°C until planting for the G2 experiment.

### Plant growth conditions

The experimental design is shown in **Figure 1**. For G1 experiment in Michigan, three germinated seeds were planted per 3.78 L (1 gallon) pot and placed into a high-light BioChambers FLEX™ LED growth chamber with a 16-hour day/8-hour night cycle at 26°C and 22°C and 50% relative humidity. When seedlings emerged and reached the earliest vegetative stage (“VC”, two cotyledons with primary leaves at nodes 1 and 2), they were thinned to one seedling per pot. Plants were watered every other day with 300 mL 0.05% 15N-10P-30K water-soluble fertilizer solution (Masterblend International, Morris, IL, USA). At the vegetative 3 stage (“V3”, third trifoliate leaves expanded), stress treatments began for the drought-treated plants. Drought plants received 100 mL of 0.15% 15N-10P-30K fertilizer solution every other day (66% less water than control but the same amount of nutrients). After approximately 14 days of treatment, when plants reached the reproductive 1 stage (“R1”, first open flowers), they were returned to the control watering every other day until senescence. There were 17 replicate plants grown per treatment in G1. Five plants per treatment were destructively harvested for plant phenotypic measurements at the reproductive 6 stage (“R6”, most pods at the seed filling stage). The remaining 12 plants were grown until senescence (S). Mature seeds were collected from the 12 senesced plants for sowing in the G2 experiment, and the root and rhizosphere microbiome were sampled from a subset of five replicate plants.

**Figure 1.**
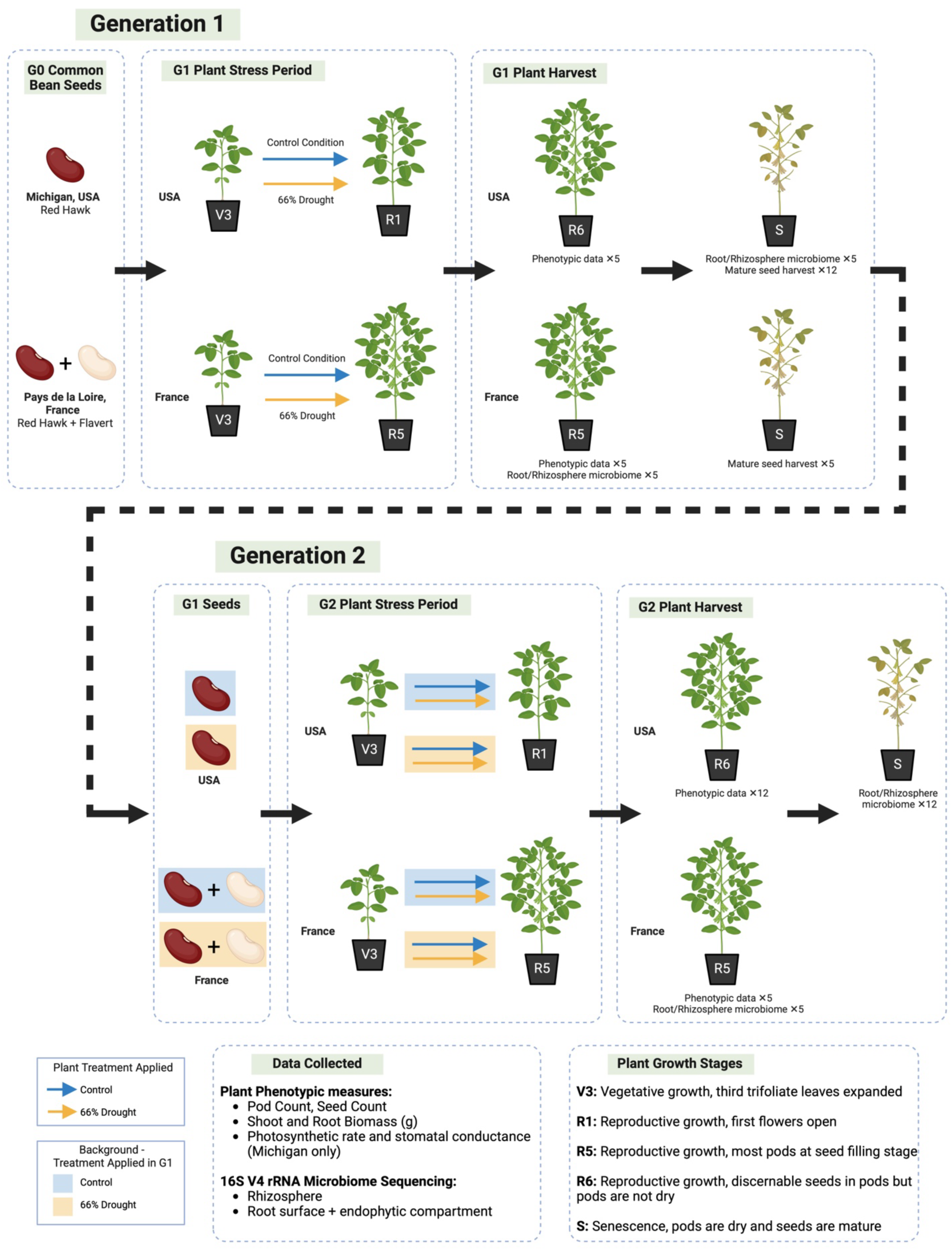
Experimental design. *Phaseolus vulgaris L.* (common bean) plants were grown over two generations in a factorial design under either control treatment with ample water and nutrients, or drought treatment with 66% less water and equal nutrient concentration to control plants. The common bean variety ‘Red Hawk’ was grown in Michigan, and both ‘Red Hawk’ and ‘Flavert’ were both grown in Pays de la Loire. Drought treatments were started at the V3 growth stage when plants had three trifoliate leaves expanded, and were concluded at the R1 stage (first open flowers) in Michigan, and at the R5 stage (half of the pods filled with discernable seeds) in Pays de la Loire. The replication indicated is per treatment, including the total number of plants used for microbiome analysis and for plant phenotyping. Image created in BioRender. Sulesky, A. (2025) https://BioRender.com/l13n679

For the G2 experiment, seeds from 24 total G1 parental lines that received either control (12 lines) or drought (12 lines) were planted. There were four combinations, indicating the G1_G2 conditions: Control_Control, Control_Drought, Drought_Control, and Drought_Drought. Seeds from each parental line were evenly distributed among the Control and Drought G2 conditions and were tracked to assess the influence of parent plant on the results. Seeds from each parental line were surface sterilized and germinated as above, then planted in the field soil-vermiculite mixture in seedling trays and placed in the growth chamber under the abovementioned conditions. When plants reached VC stage, four viable seedlings per parent line were transferred to 3.78 L (1-gallon) pots for the remainder of the experiment. Thus, each parental line had two offspring directed into each of the G2 conditions. One offspring was used for plant phenotypic measurements (12 plants total per G2 condition), while the other was harvested for microbiome analysis (12 plants total per G2 condition). Plants were watered according to the timeline described for G1.

Pays de la Loire plants were grown under the same growth chamber, control, and drought conditions above. The Pays de la Loire experiment was replicated with Red Hawk and Flavert seeds. After three weeks of growth, day 18 after sowing (V3 stage), replicate plants (n= 5) were exposed to control conditions (300 mL of 0.05% nutritive solution) or drought (66% water-withholding, 100 mL of 0.15% nutritive solution) for four weeks until Day 56 within the R5, which was a slightly longer stress period than applied to the plants in Michigan. Five plant parental lines from each G1 condition (10 lines total) from both genotypes were planted in G2. They were also planted in a complete factorial design with four treatment conditions in G2.

### Plant phenotypic data

Plants dedicated to phenotypic trait measurements at Michigan State were collected with a LI-COR LI-6800 to measure photosynthetic rate and stomatal conductance on the day before the drought ended within stage R1 (LI-COR Biosciences, Lincoln, NE, USA). Plants were grown until they reached approximately the reproductive 6 stage (“R6”, pods developed with discernable green seeds that were not yet dry). Pods were removed and placed in a paper bag, plants were cut at the base of the stem, and shoots were placed in a large envelope. Roots were removed from the pot, shaken to remove excess soil, collected, rinsed, and put in a paper bag. Pods and seeds per plant were counted, and then all three compartments were placed in a 50 °C drying oven for two days. After drying, the shoot, root, and pod dry biomasses were measured.

In Pays de la Loire, plants were harvested for trait measurements at approximately the R5 stage. The entire root system was gently separated from the soil in the pot and placed in a plastic bag. The total fresh weight of each plant (both above- and belowground tissues) was measured, and the number of pods and seeds per plant was counted.

### Microbiome compartment harvest

Once the Michigan plants had senesced and pods were dried, plants were harvested for microbiome analysis. Seed pods were removed and stored in a sterile Whirl-pak® bag. Root systems were removed from the pot and shaken to remove loose soil. Roots were collected in a Whirl-pak® bag, and associated rhizosphere soil was collected in a sterile 50 mL conical tube. Roots and rhizosphere soil were stored at −80°C until nucleic acids extraction.

The microbiome analysis in Pays de la Loire was performed on plants that were harvested at approximately the R5 stage. The entire root system was gently separated from the soil in the pot and placed in a plastic bag. The rhizosphere soil was collected by shaking the root system in a plastic bag and then it was stored at −80°C. The root system was washed in sterile distilled water, transferred to 50 mL tubes, and stored at −80°C.

### DNA extractions

The Michigan root samples were thawed at room temperature, and a 3 to 6 cm section of the main root system was cut and used for root DNA extraction, which combined the rhizoplane and endophytic bacteria of the root tissues. The selected sections were rinsed with sterile DI water, and finely ground in liquid nitrogen with a mortar and pestle. DNA was extracted from the ground root material with the DNeasy PowerSoil Pro DNA Kit (Qiagen, Germantown, MD, USA) following the manufacturer’s instructions with the following modifications. In step one, 750 μL solution CD1 was used with 50 μL ATL buffer (Qiagen, Germantown, MD, USA). The bead beating step was performed for 15 minutes on a vortex genie 24-tube adapter at maximum speed. Lastly, 60 μL of the final elution buffer C6 was used, and tubes were incubated for 10 minutes before centrifugation.

Rhizosphere soil was thawed at room temperature, and DNA was extracted using the DNeasy PowerSoil Pro DNA Kit with the same ATL buffer modification as described for the roots. Negative controls (extraction reagent blanks) were included with each batch of DNA extractions, and one positive mock community control was included with each compartment sample set (roots or rhizosphere) (Colovas *et al*. 2022). Controls were sequenced alongside the experimental samples.

French samples were processed similarly and extracted using the DNeasy PowerSoil Kit (Qiagen, Germantown, MD, USA, *discontinued*) following the manufacturer’s instructions. A blank extraction kit control, a PCR-negative control, and a PCR-positive control (*Lactococcus piscium* DSM 6634, a fish pathogen that is not plant-associated) were included in each PCR plate.

### Sequencing

The 16S V4 rRNA gene was sequenced for the Michigan root and rhizosphere samples at the Argonne National Laboratory Environmental Sample Preparation and Sequencing Facility (Lemont, IL, USA). The DNA was PCR amplified with region-specific primers that included sequencer adapter sequences used in the Illumina MiSeq; FWD: GTGYCAGCMGCCGCGGTAA; REV: GGACTACNVGGGTWTCTAAT (Caporaso *et al*. 2011, 2012; Apprill *et al*. 2015; Walters *et al*. 2015; Parada, Needham and Fuhrman 2016). Each 25 µL PCR reaction contained 9.5 µL of MO BIO PCR Water (Certified DNA-Free), 12.5 µL of QuantaBio’s AccuStart II PCR ToughMix (2x concentration, 1x final), 1 µL Golay barcode tagged Forward Primer (5 µM concentration, 200 pM final), 1 µL Reverse Primer (5 µM concentration, 200 pM final), and 1 µL of template DNA. The conditions for PCR were as follows: 94 °C for 3 minutes to denature the DNA, with 35 cycles at 94 °C for 45 s, 50 °C for 60 s, and 72 °C for 90 s, with a final extension of 10 min at 72 °C to ensure complete amplification. Amplicons were then quantified using PicoGreen (Invitrogen) and a plate reader (InfiniteÒ 200 PRO, Tecan). Once quantified, the volumes of each of the products were pooled into a single tube in equimolar amounts. This pool was then cleaned using AMPure XP Beads (Beckman Coulter) and quantified using a fluorometer (Qubit, Invitrogen). After quantification, the pool was diluted to 2 nM, denatured, and then diluted to a final concentration of 6.75 pM with a 10% PhiX spike for sequencing. Amplicons were sequenced on a 251bp x 12bp x 251bp MiSeq run using customized sequencing primers and procedures (Caporaso *et al*. 2012).

For the 16S V4 rRNA gene sequencing in Pays de la Loire, PCR reactions were performed with a high-fidelity Taq DNA polymerase (AccuPrimeTM Taq DNA Polymerase System, Invitrogen) using 5μL of 10X Buffer, 1μL of forward and reverse primers (10μM), 0,2μL of Taq and 10μL of DNA. A first PCR amplification was performed with the primer sets V4 515f/806r (5‘-GTGCCAGCMGCCGCGGTAA-3’and 5’-GGACTACHVGGGTWTCTAAT-3’ (Caporaso *et al*. 2011). Cycling conditions were composed with an initial denaturation at 94°C for 3 minutes followed by 35 cycles of denaturation at 94°C (30 seconds), primer annealing at 55°C (45 seconds), and extension at 68°C (90 seconds), with a final elongation at 68°C for 10 minutes. Amplicon purification was performed with a ratio of 0.8 of magnetic beads (Sera-MagTM, Merck). A second PCR amplification was performed to incorporate Illumina adapters and barcodes: a first denaturation at 94°C (1 minute), followed by 12 cycles of denaturation at 94°C (60 seconds), primer annealing at 55°C (60 seconds) and extension at 68°C (60 seconds) with a final elongation at 68°C for 10 minutes. Amplicons were purified with a ratio of 0.7 of magnetic beads and quantified with the Quant-iTTM PicoGreen® dsDNA Assay Kit (Invitrogen). All the amplicons were pooled in equimolar concentrations, and the concentration of the equimolar pool was monitored with quantitative PCR (KAPA SYBR® FAST, Merck). Amplicon libraries were mixed with 10% PhiX and sequences with a MiSeq reagent kit v3 600 cycles (Illumina).

### Sequence data processing

Fastq files were processed in QIIME2 (v.2022.8.0) after primer removal (Bolyen *et al*. 2019) and demultiplexed with the demux emp-paired protocol. Samples were denoised, truncated, and merged at 100% sequence identity using DADA2 (Callahan *et al*. 2016) in QIIME2, with the truncation lengths found in supplemental Table S1. 16S rRNA gene taxonomy was assigned with the SILVA database (Quast *et al*. 2013) release 132 for French datasets and release 138 for Michigan datasets, and taxonomy and amplicon sequence variants (ASV) tables were exported for further analysis in R.

### Plant phenotypic and microbiome data analysis

Data analyses were performed in R v.4.4.1 (R Core Team 2024) and R Studio v.2025.5.1.513 (Posit team 2025). Amplicon sequence variants, taxonomy, and metadata tables were imported into the phyloseq package v.1.48.0 (McMurdie and Holmes 2013). Sequences derived from 16S rRNA genes unclassified at the phylum level or affiliated with Chloroplasts or Mitochondria were removed. The identification of sequence contaminants was assessed with decontam package v.1.20.0 (Davis *et al*. 2018) using the prevalence of ASVs in samples and negative controls (Table S1). Rarefaction curves for each dataset were generated using the rarecurve() function in the vegan package v.2.6.4 (Oksanen *et al*. 2020), and multiple rarefactions (without replacement, iterations of 100) were performed for each datasets using the phyloseq_mult_raref () function in the metagMisc package v.0.5.0 (Mikryukov 2025). The average of observed richness was calculated across multiple rarefactions using the phyloseq_mult_raref_div() function in the metagMisc package. The effects of drought treatment (G1 and G2) and plant genotype, as well as their interactions on richness were assessed with Welch’s two sample t-test. Effect of drought treatment and plant genotype and their interaction was assessed by two-way ANOVA. In addition, a three-way ANOVA was performed to analyse the effect of drought, genotype and plant generation. Tukey Honest Significant Difference (Tukey’s HSD) post-hoc test were performed where applicable. The normality of residuals and homogeneity of the residual variances were verified using Shapiro-Wilk and Levene’s tests, respectively. Response variable data were transformed when necessary using arcsinh or box-cox or ordered quantile normalization (orderNorm) implemented in the bestNormalize package v.1.9.1 (Peterson and Cavanaugh 2020; Peterson 2021). Microbiome alpha diversity figures were created using the ggplot2 package v.3.5.2 (Wickham 2016).

Bray-Curtis distances were calculated and averaged across rarefactions using the mult_dissim() and mult_dist_average() functions in the MetagMisc package. Permutational multivariate analysis of variance (PERMANOVA) was performed with the adonis2() function with 9999 permutations from the vegan package to assess the effect of drought treatment, plant genotype, and their interactions on microbiome community composition. Due to the significant effect of planting group in the G2 Michigan data, we restricted the permutation by planting group using the setBlock() function in the permute package v.0.9.8 (Simpson 2025). Post-hoc analysis on the PERMANOVA results was performed with the pairwise.adonis2() function from the pairwiseAdonis package v.0.4.1 (Arbizu 2017). Principal coordinates analysis (PCoA) plots were created from Bray-Curtis dissimilarities using the cmdscale() function in the stats package v.4.4.1 with ggplot2 for visualization. For the G2 Michigan data, we performed partial constrained analysis of principal coordinates using capscale() with Condition() function in the vegan package to control the effect of planting group. Beta dispersion was assessed with the betadisper() and permutest() functions from the vegan package with the spatial median method (Anderson 2006).

The effect of drought, plant genotype, and their interactions on plant phenotypes were analyzed with Welch’s two sample t-test or two-way ANOVA or three-way ANOVA with Tukey’s HSD post-hoc tests where applicable. The data normality and homogeneity were assessed using Shapiro-Wilk and Levene’s tests, respectively. Response variable data were arcsinh or square-root transformed as needed using the bestNormalize package. Plant phenotype figures were created with ggplot2. Additional data processing and statistics were performed in the tidyverse package v.2.0.0 (Wickham *et al*. 2019) and R stats. Figure panels were assembled with the patchwork package v.1.3.0 (Pedersen 2024).

### Data Availability

Data analysis code can be found at (https://github.com/ShadeLab/Drought_multigeneration_study_common_bean). Raw sequence data from the French samples can be found on the European Nucleotide Archive under accession number PRJEB65346. Raw sequences from the Michigan samples can be found on the NCBI Sequence Read Archive under BioProject accession number PRJNA1058980.

## Results

Because the two locations had several necessarily differing parameters (see methods), differences in their microbiomes were expected. The datasets from Michigan and Pays de la Loire were quality controlled and analyzed independently, and the comparative analyses were focused on overall plant and microbiome trends and dynamics. Our objective was to assess ecological patterns and generalities in consecutive, multi-generational drought response in the microbiome despite differences due to different soils and geographic locations.

### Plants responded differently to drought in different locations

Plant biomass, yield, and photosynthetic data were collected. For plants grown in Pays de la Loire, France, with a slightly longer stress period, both Red Hawk and Flavert had a decrease in the number of pods on the drought-treated plants (Welch Two Sample t-test, Red Hawk: t=4.26, P= 0.004; Flavert: t=7.06, P= 0.0001) (Fig. 2, A.1, Table S2, S3). However, the decrease in pod number did not correspond to a decrease in seed number (Fig. 2, A.2). For Red Hawk beans grown in Michigan, the numbers of pods, seeds, and root biomass were not affected by drought (Fig. 2, B.3, B.4, B.6, Table S2). However, the photosynthetic rate (Welch Two Sample t-test: t=3.28, P=0.01), stomatal conductance (Welch Two Sample t-test: t=2.82, P=0.02), and above-ground mass (Welch Two Sample t-test: t=2.49, P=0.03) were reduced in the drought plants as compared to the control plants (Fig. 2, B.1, B.2, B.5, Table S2). These results generally indicate that there was a negative effect of drought on the G1 plants, but that the precise effect was inconsistent across the different locations.

**Figure 2.**
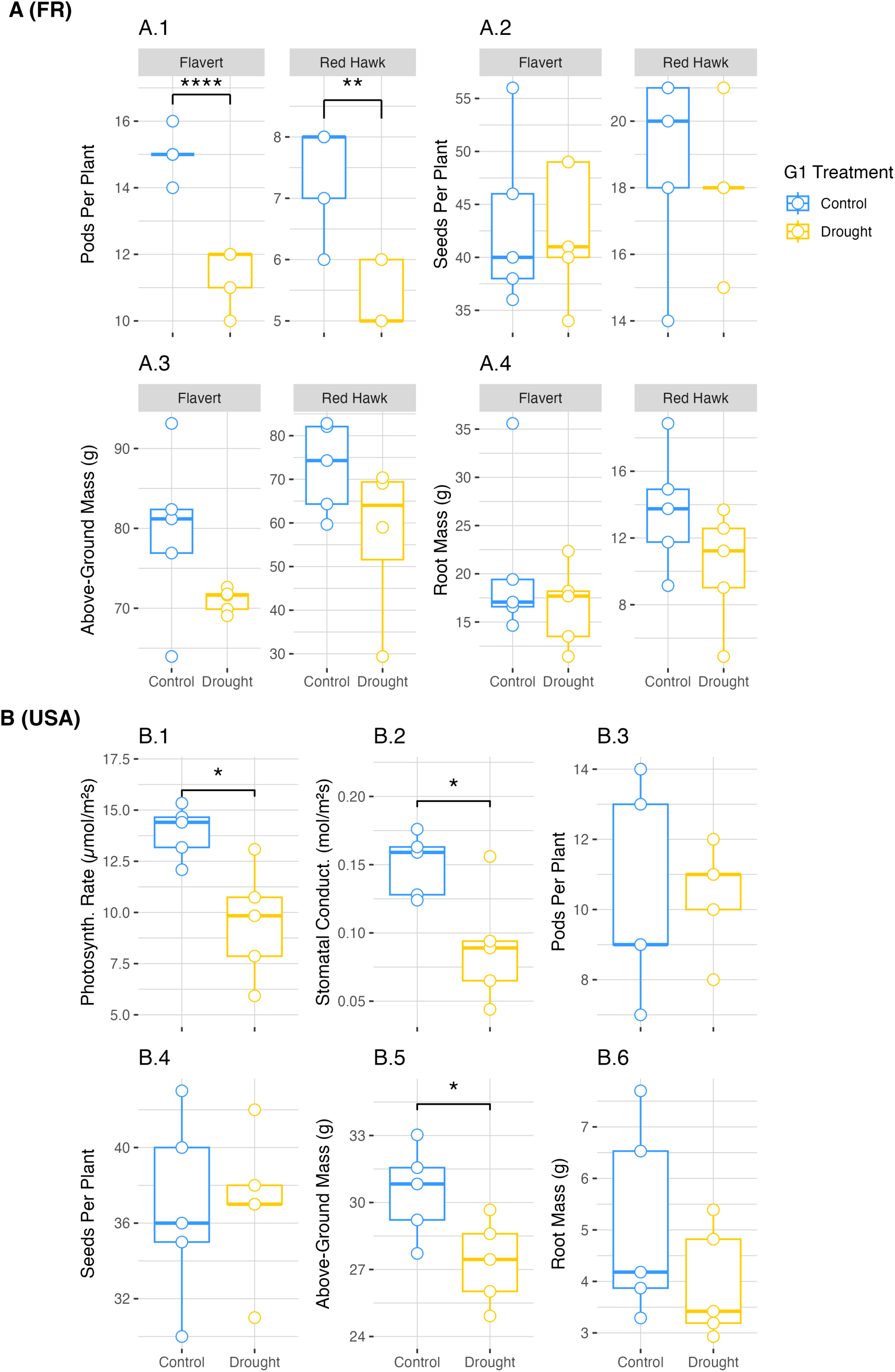
Plant phenotypic measurements in G1. **A.** Phenotypic measurements taken for common bean plants of both Flavert and Red Hawk genotypes grown in Pays de la Loire, France. Above-ground and root biomass measurements were taken on fresh plant tissue. **B.** Phenotypic measurements were taken for common bean Red Hawk plants grown in Michigan, USA. Above-ground and root biomass measurements were taken on dry plant tissue. All above-ground biomass measurements include the total mass of stems, leaves, and pods. Photosynth. Rate = Photosynthetic Rate, Stomatal Conduct. = Stomatal Conductance. Two-way ANOVA with Tukey’s HSD post hoc test (Pays de la Loire, France) and Welch Two Sample t-test (Michigan, USA), * = p-value < 0.05, ** = p-value < 0.01, **** = p-value < 1e-4, n=5 plants per treatment.

There was a strong effect of G2 drought on the Flavert (two-way ANOVA, pod count: F=85.63, P=7.99e-08; seed count: F=85.79, P=7.89e-08, shoot mass: F=161.86, P=8.78e-10) and Red Hawk (two-way ANOVA, pod count: F=28.13, P=7.14e-05; seed count: F=141.75, P=2.31e-09, shoot mass: F=225.98, P=7.4e-11; root mass: F=21.95, P=0.0002) plants grown in Pays de la Loire (Table S4, S5). However, the observed G2 drought effect was independent of the G1 condition. Specifically, the number of pods, seeds, and above-ground biomass were decreased for both bean genotypes grown in Pays de la Loire (Fig. 3, A). In contrast, the root mass of Red Hawk in Pays de la Loire was increased by drought (Fig. 3, A.4). For Red Hawk plants grown in Michigan, only the photosynthetic rate (two-way ANOVA: F=18.43,P=0.0006) and stomatal conductance (two-way ANOVA: F=46.8,P= 5.59e-06) were decreased by drought, like what was observed during the G1 drought (Fig. 3, B.1, B.2, Table S4). These data indicate that the Flavert and Red Hawk plants in Pays de la Loire were more negatively impacted by the drought in G2 than the plants in Michigan. The legacy of G1 condition did not affect most plant outcomes in G2, except for Red Hawk biomass in Pays de la Loire. Flavert plants that were droughted in both generations had lower above-ground biomass than plants that were not droughted in G1 (Fig. 3, A.3).

**Figure 3.**
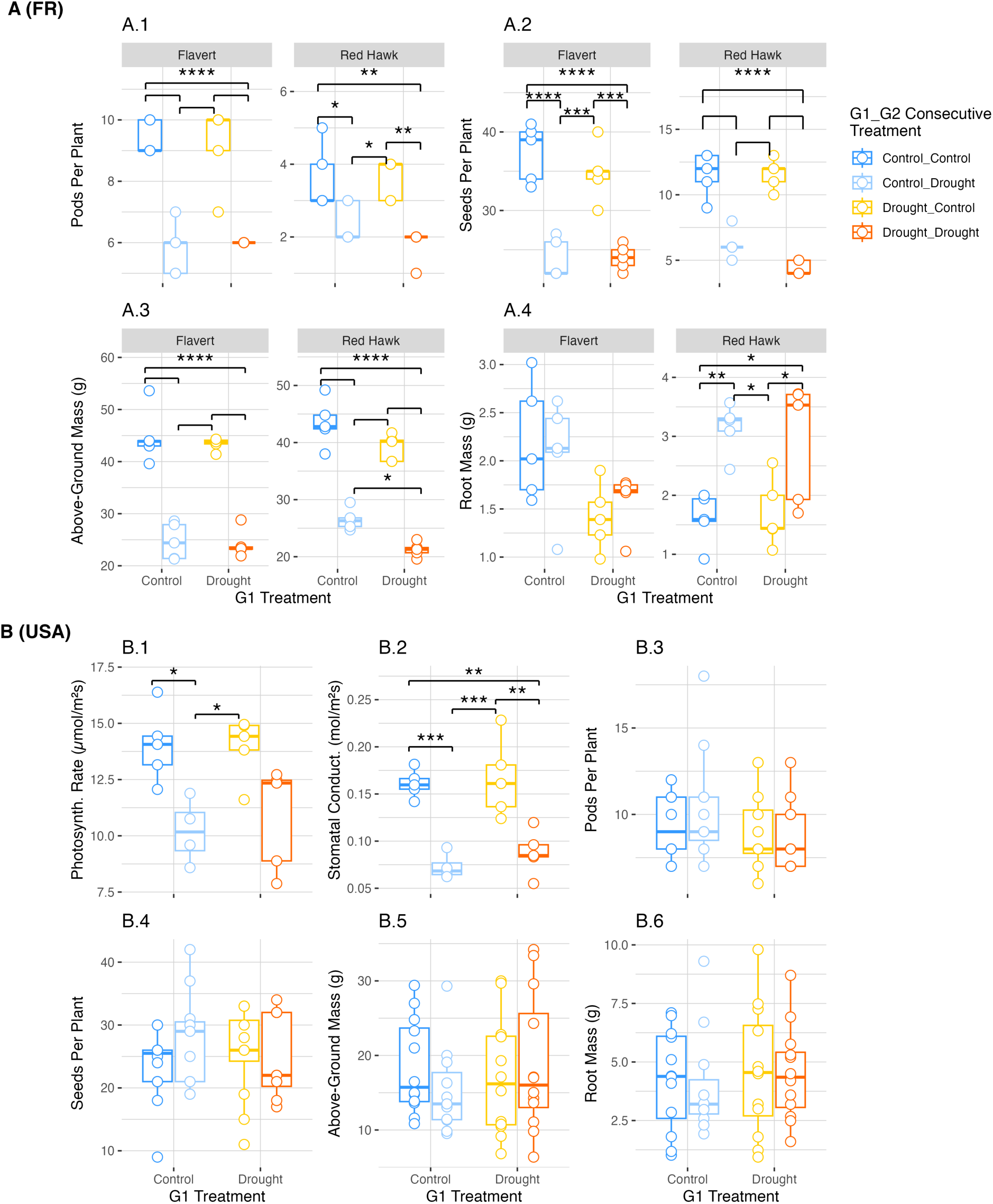
Plant phenotype measurements in Generation 2. **A.** Phenotypic measurements taken for common bean plants of both Flavert and Red Hawk genotypes grown in Pays de la Loire, France. Pays de la Loire had n=5 plants per treatment. **B.** Phenotypic measurements taken for common bean Red Hawk plants grown in Michigan, USA. Michigan had n=12 plants per treatment. Above-ground and root biomass measurements were taken on dry plant tissue. Above-ground biomass measurements include the total mass of stems, leaves, and pods. Photosynth. Rate = Photosynthetic Rate, Stomatal Conduct. = Stomatal Conductance. Two-way ANOVA with post-hoc Tukey HSD test, * = p-value < 0.05, ** = p-value < 0.01, *** = p-value < 0.001, **** = p-value < 1e-4. Non-annotated significance lines have the same p-value as lines above.

### Bacterial community alpha diversity is decreased by drought in Flavert plants

Overall, the bacterial community richness (number of observed ASVs) in Pays de la Loire microbiomes was lower than the alpha diversity observed in Michigan samples. Sequencing coverage was sufficient for both datasets, as the rarefaction curves reached asymptotes (Fig. S1). There were differences in richness between the two genotypes grown in Pays de la Loire in root (two-way ANOVA: F=7.651, P=0.0138; Tukey’s HSD: P.adj<0.05) but not in rhizosphere samples of drought-stressed plants in G1 (Fig. 4A, C, Table S6). However, these differences were not present in control plants. Global analysis revealed no effect of drought in G1 in both plant compartments in Pays de la Loire (Table S6). Similarly, the G1 microbiome alpha diversity of Red Hawk in Michigan was unaffected by drought (Welch Two Sample t-test, root: t=0.055, P=0.958; rhizosphere: t=0.792, P=0.472) (Fig. 4 B, D, Table S6).

**Figure 4.**
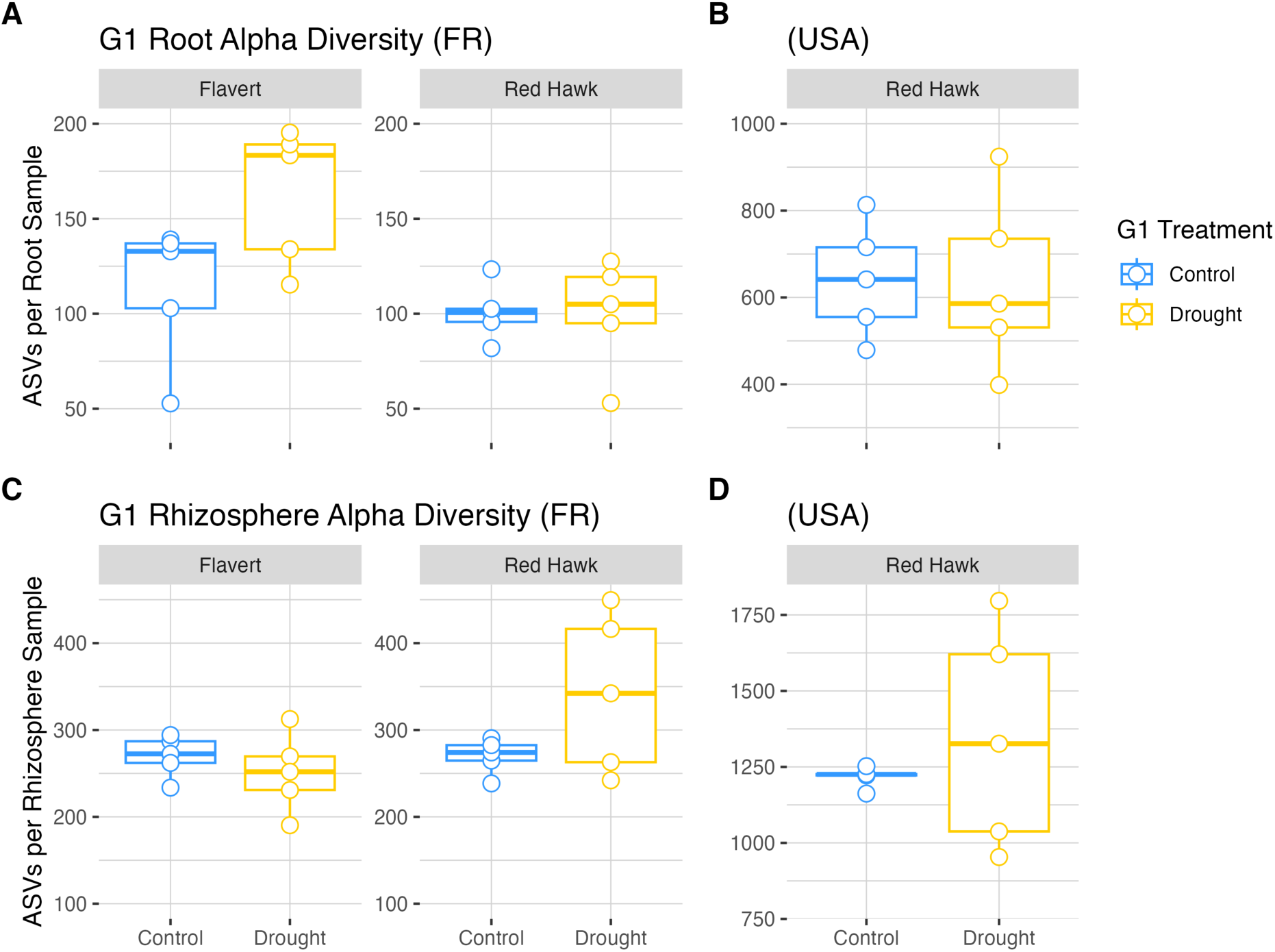
Alpha diversity observed for root and rhizosphere samples in Generation 1. **A.** Root microbiomes of plants grown in Pays de la Loire, France. **B.** Root samples of plants grown in Michigan, USA. **C.** Rhizosphere samples of plants grown in France. **D.** Rhizosphere samples of plants grown in the USA. Plant genotype is indicated by bars above the panels. Two-way ANOVA with Tukey’s HSD post hoc test (Pays de la Loire, France) and Welch Two Sample t-test (Michigan, USA) (n=5 per treatment). Absence of annotation indicates no significant difference (p-value > 0.05).

In the G2 experiment, we observed an effect of G2 drought on root microbiome alpha diversity in Pays de la Loire (three-way ANOVA: F=7.023, P=0.0135), but there was no legacy of G1 drought (three-way ANOVA, genotype x G1x G2: F=0.03, P=0.86) (Table S7). Further *post-hoc* analysis showed no differences between groups for Flavert root microbiomes in Pays de la Loire, indicating the drought effect was marginal (Fig. 5A). There also was no legacy effect of drought detected for Michigan Red Hawk root microbiomes (two-way ANOVA, G1 x G2: F=1.45, P=0.23) (Fig. 5B, Table S7). The two genotypes grown in Pays de la Loire were different in their rhizosphere microbiome alpha diversity (three-way ANOVA: F=4.983, P=0.03378; Tukey’s HSD: P.adj<0.001) (Fig. 5C, Table S7). We observed a significant effect of G1 drought (three-way ANOVA: F=9.132, P=0.00532) on the richness of rhizosphere microbiome in Pays de la Loire (Table S7). Interestingly, the effects of drought were more pronounced in Flavert G2 plants that experienced drought in G1 with a greater decrease in richness compared to G1 control plants (Tukey’s HSD, Control_Control v. Drought_Control: P.adj<0.01, Control_Control v. Drought_Drought: P.adj<0.01) (Fig. 5C, Table S8).

**Figure 5.**
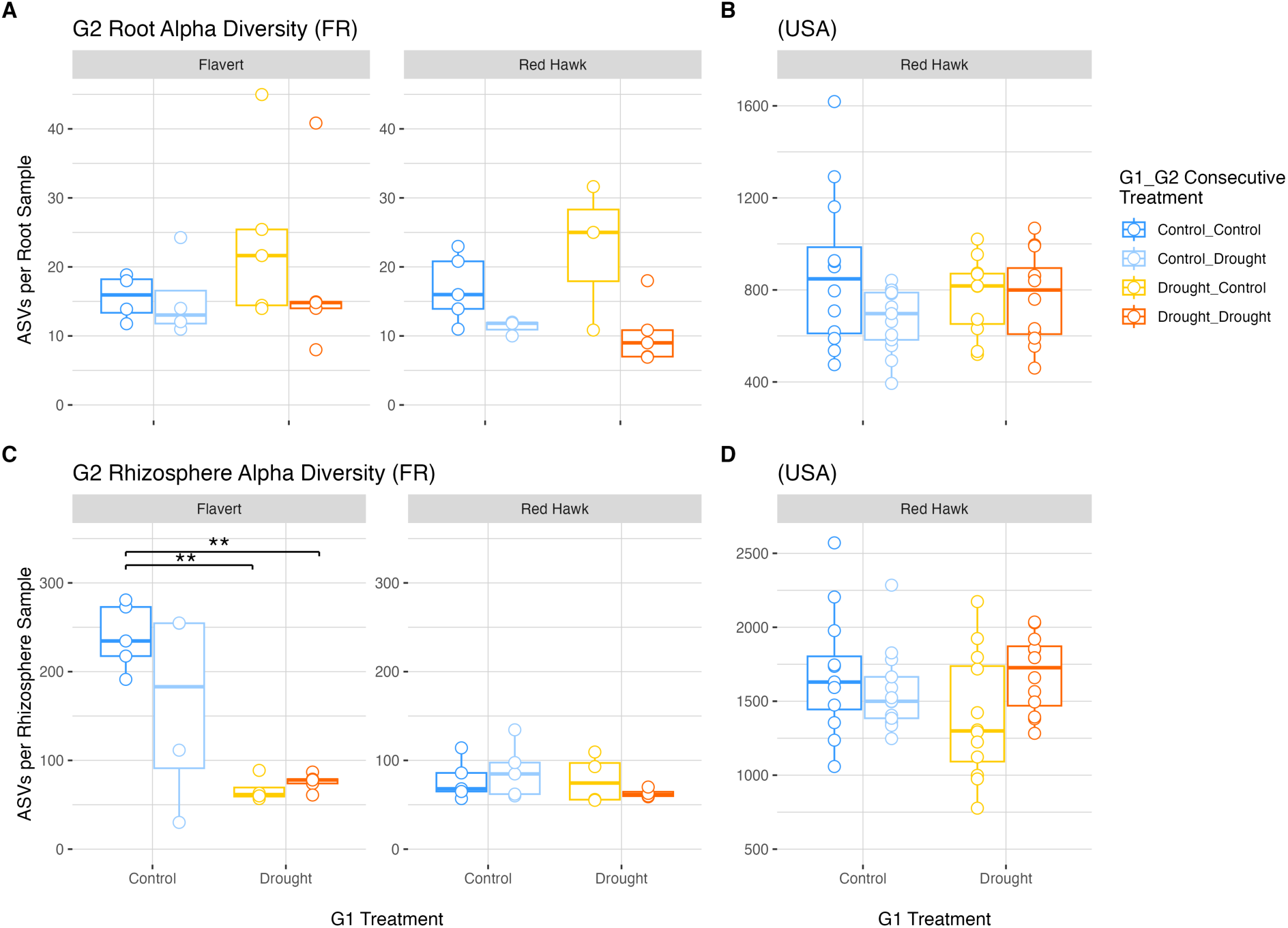
Alpha diversity observed for root and rhizosphere samples in Generation 2. **A.** Root samples of plants grown in Pays de la Loire, France (n=5 plants per treatment). **B.** Root samples of plants grown in Michigan, USA (n = 12 plants per treatment). **C.** Rhizosphere samples of plants grown in France. **D.** Rhizosphere samples of plants grown in the USA. Plant genotype is indicated by bars above the panels. Two-way ANOVA with Tukey’s Honest Significant Difference test, ** = p-value < 0.01.

Overall, genotype was found to play a significant role in determining root and rhizosphere alpha diversity, and Flavert plants had decreased alpha diversity under drought conditions in G2, with no legacy effect detected. A global interaction between G1 and G2 treatments was initially detected in the rhizosphere microbiome of Red Hawk in Michigan (two-way ANOVA, G1 x G2: F=4.145, P=0.047) (Table S7), but it was not supported by *post-hoc* pairwise tests. Thus, a legacy of the G1 drought on the G2 outcome was not reflected in Red Hawk rhizosphere microbiome in either location, and, furthermore, the Red Hawk rhizosphere alpha diversity was not affected by the drought in G2 (Tukey’s HSD: P.adj>0.05) (Fig. 5C, D). Thus, despite that Red Hawk plant traits were impacted by drought in Pays de la Loire, these plants’ alpha diversity was unaffected.

### Inconsistent drought and drought legacy effects on bacterial community structure

Beta diversity of the root and rhizosphere microbial communities were analyzed independently for Pays de la Loire and Michigan plants. In G1, there were differences in the bacterial community structure of the two genotypes grown in Pays de la Loire in root (PERMANOVA: R^2^=0.49, F=17.09, P=0.0001) (Fig. 6A, B, Table S9), but not in rhizosphere (PERMANOVA: R^2^=0.06, F=1.22, P=0.22) (Fig. 6D, E) samples. When the genotypes were analyzed separately, the drought treatment in G1 had no influence on the root beta diversity in Pays de la Loire (PERMANOVA, Red Hawk: R^2^=0.11, F=1.01, P=0.39; Flavert: R^2^=0.12, F=1.05, P=0.35) (Table S10). Drought mildly affected the beta diversity in G1 rhizosphere (PERMANOVA: R^2^=0.09, F=1.72, P=0.02), but not in root (PERMANOVA: R^2^=0.03, F=1.07, P=0.31) (Table S9). We also found that the effect of drought on the rhizosphere communities in G1 differed by genotype. For example, when we analyzed separately, there were significant differences between drought and control in the rhizosphere beta diversity for Flavert (PERMANOVA: R^2^=0.16, F=1.58, P=0.02) (Fig. 6D, Table S10), but not for Red Hawk (PERMANOVA: R^2^=0.13, F=1.22, P=0.1) (Fig. 6E, Table S10) genotype. However, the drought treatment did not affect the root or rhizosphere beta diversity in Michigan plants in G1 (PERMANOVA, root: R^2^=0.15, F=1.43, P=0.14; rhizosphere: R^2^=0.1, F=0.97, P=0.43) (Fig. 6C, F, Table S9).

**Figure 6.**
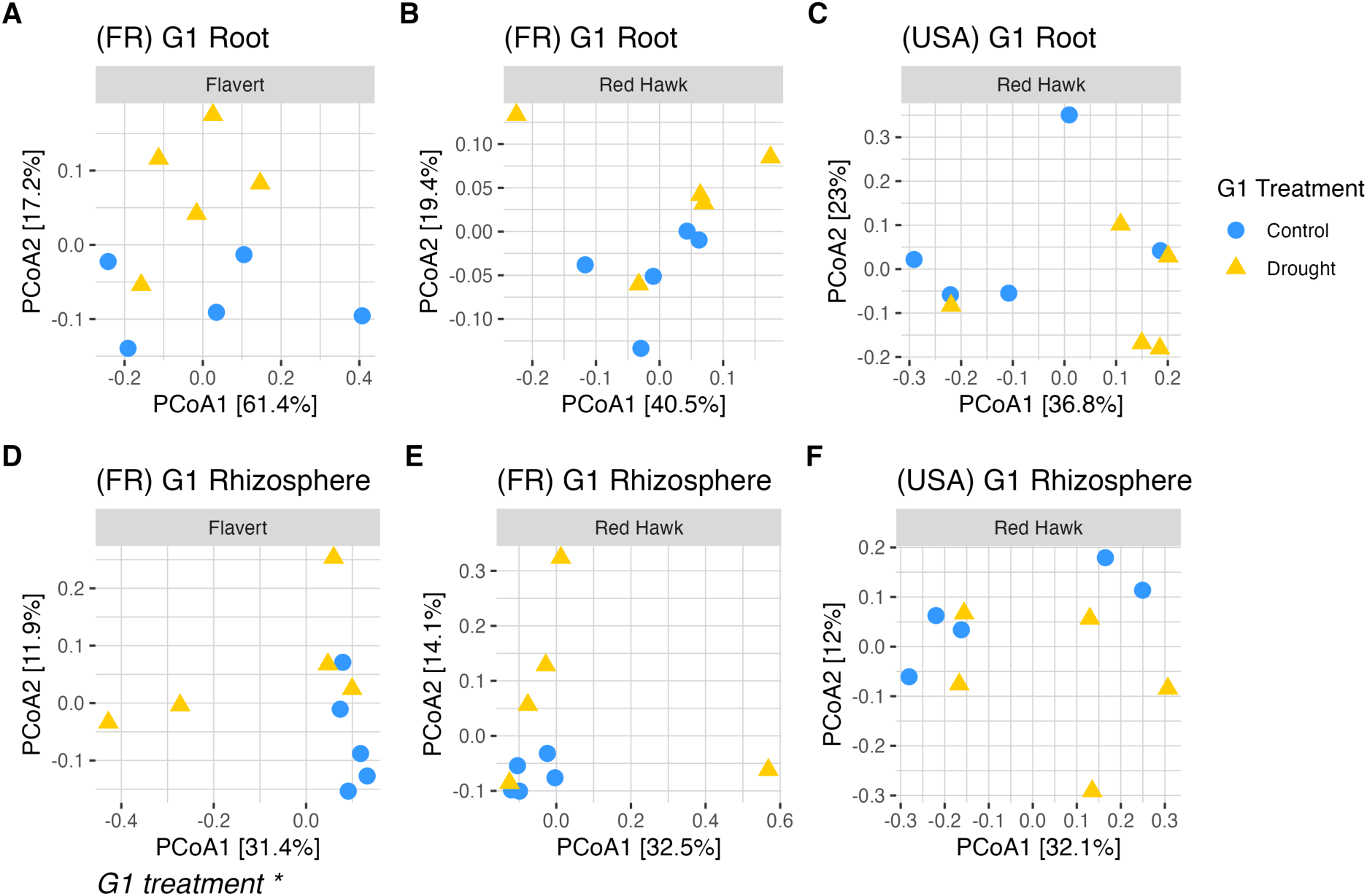
PCoA ordinations of beta diversity Bray-Curtis dissimilarities in the Generation 1 (G1) experiment. **A, B.** Root samples of Flavert and Red Hawk plants grown in Pays de la Loire, France. **C.** Root samples of plants grown in Michigan, USA. **D, E.** Rhizosphere samples of Flavert and Red Hawk plants grown in Pays de la Loire, France. **F.** Rhizosphere samples of plants grown in Michigan, USA. There were significant differences between genotypes in root sample in France (PERMANOVA: R^2^=0.49, F=17.09, P=0.0001). Plant genotype is indicated by the panel labels. * = p-value < 0.05, n=5 per treatment.

In G2, there were also differences between the two plant genotypes in Pays de la Loire for both compartments (PERMANOVA, root: R^2^=0.31, F=15.47, P=0.0006 (Fig. 7A, B, Table S11); rhizosphere: R^2^=0.06, F=2.57, P=0.0007 (Fig. 7D, E, Table S11)). Flavert root beta diversity was affected by the interaction of the G1 x G2 treatments (PERMANOVA: R^2^=0.25, F=4.94, P=0.02) (Fig. 7A, Table S12), indicating that the effect of drought in the G2 was affected by its G1 treatment. Pairwise multilevel comparisons of Flavert’s root microbiome showed distinct community structures between the Control_Drought and Drought_Drought plants (PERMANOVA: R^2^=0.42, F=5.1, P=0.018). Red Hawk roots in G2 in Pays de la Loire were unaffected by G1 drought (PERMANOVA: R^2^=0.03, F=0.41, P=0.59) and G2 drought (PERMANOVA: R^2^=0.06, F=0.86, P=0.38) (Fig. 7B, Table S12). However, a mild effect of G2 drought was observed on Red Hawk roots in Michigan (PERMANOVA: R^2^=0.03, F=1.25, P=0.029) (Fig. 7C, Table S11).

**Figure 7.**
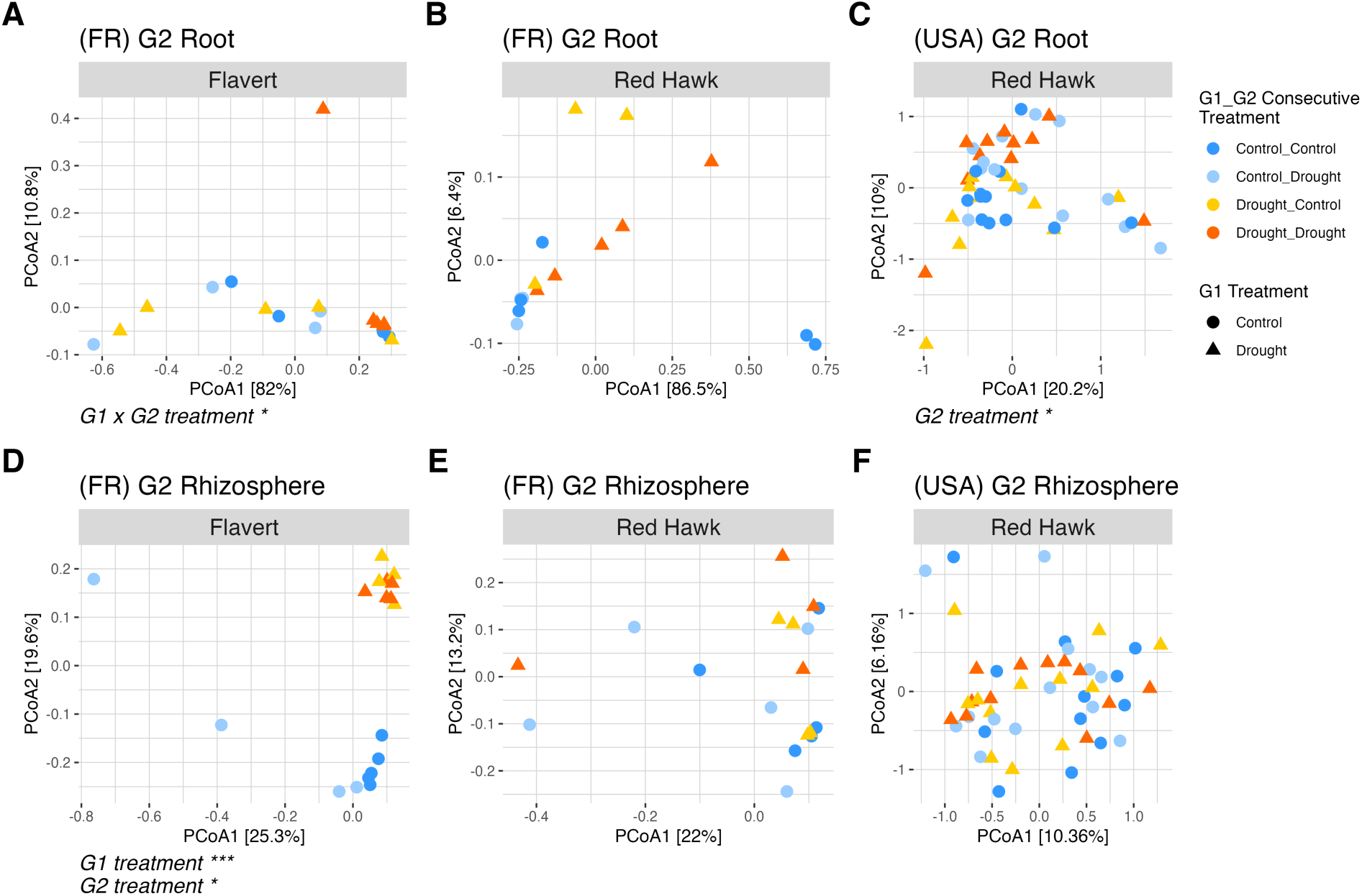
PCoA ordinations of Bray-Curtis dissimilarities in the Generation 2 experiment (G2). **A, B.** Root samples of Flavert and Red Hawk plants grown in Pays de la Loire, France. **C.** Root samples of plants grown in Michigan, USA. **D, E.** Rhizosphere samples of Flavert and Red Hawk plants grown in France. **F.** Rhizosphere samples of plants grown in the USA. There were differences between genotypes in both root (PERMANOVA: R^2^=0.31, F=15.47, P=0.0006) and rhizosphere (PERMANOVA: R^2^=0.06, F=2.57, P=0.0007) samples in France. Plant genotype is indicated by panels labels * = p-value < 0.05, *** = p-value < 0.001. France n=5 plants per treatment, USA n=12 plants per treatment.

In the G2 rhizosphere samples from Pays de la Loire, the impact of drought was influenced by genotype, as demonstrated by a significant interaction between genotype and G1 treatment (PERMANOVA: R^2^=0.07, F=2.75, P=0.0006) (Table S11). Flavert plants exhibited distinct rhizosphere communities based on G1 legacy condition (PERMANOVA: R^2^=0.2, F=4.32, P=0.0001) (Fig. 7D, Table S12), while Red Hawk plants were not affected by either G1 or G2 condition. In comparison, Red Hawk rhizosphere communities in Michigan were not affected by either generation (PERMANOVA, G1xG2: R^2^=0.017, F=0.81, P=0.18) or condition (PERMANOVA, G2: R^2^=0.015, F=0.73, P=0.39) (Fig. 7F, Table S11).

Often, disturbed systems exhibit relative increases in their variability (Zaneveld, McMinds and Vega Thurber 2017). Thus, we also analyzed the dispersion of the beta diversity (beta dispersion) within and between growth conditions and generations. In G1 plants, a minor effect of drought was only observed in rhizosphere communities of Flavert in Pays de la Loire with increased dispersion (PERMDISP: F=4, P=0.046) (Fig. 8A, Table S13). In G2, root communities of Flavert plants in Pays de la Loire was unaffected by drought (PERMDISP: P>0.05) (Table S14). Meanwhile, we found that rhizosphere communities of Flavert G2 in Pays de la Loire had altered beta dispersion (PERMDISP, G2: F=4.05, P=0.048; G1 x G2: F=4.12, P=0.02) (Table S14), with Control_Drought plants having higher dispersion than Control_Control in the root samples (Pairwise PERMDISP: P<0.05) (Fig. 8C). Beta dispersion in Red Hawk G2 plants was unaffected by drought in Pays de la Loire, despite there was a difference between Control_Drought and Drought_Control plants (Pairwise PERMDISP: P<0.05, Fig. 8B). Red Hawk plants in Michigan had lower dispersion in the rhizosphere of droughted plants in G2, regardless of G1 treatment (Pairwise PERMDISP: P<0.05). Consistent with the effects observed in alpha and beta diversity, the dispersion of the bacterial rhizosphere community in the Control_Drought Flavert plants was altered in G2, indicating legacy effects for these plants. The Red Hawk rhizosphere beta dispersion was affected by the G2 drought in Michigan (Fig. 8D).

**Figure 8.**
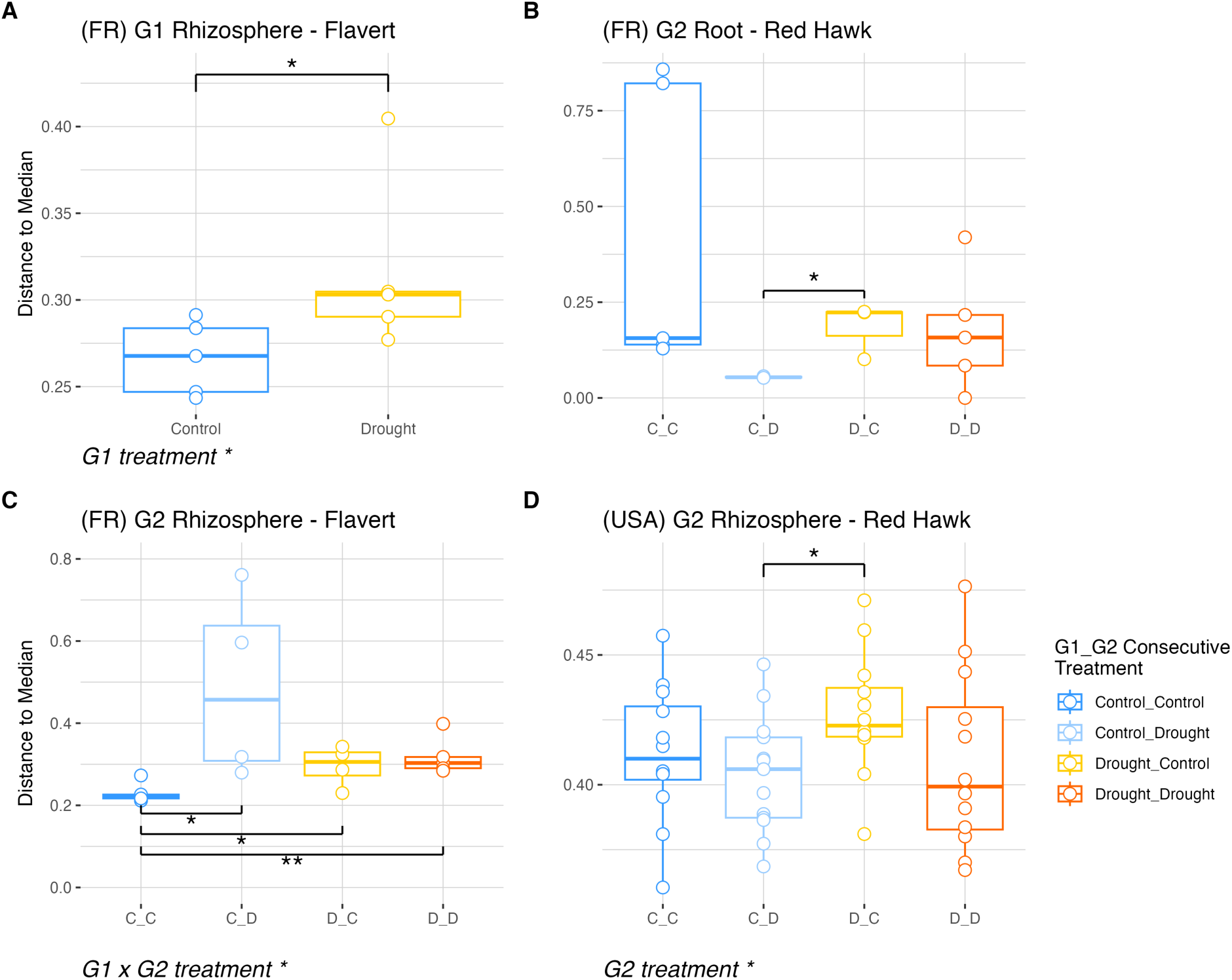
Beta dispersion calculated on the spatial median between samples. Only figures showing groups with statistically supported differences are shown. All other ordinations had no significant differences in dispersion between groups. **A.** Distance to the median in G1 Flavert rhizosphere in Pays de la Loire, France. **B.** Distance to the median in G2 Red Hawk roots in Pays de la Loire, France. **C.** Distance to median in G2 Flavert rhizosphere in Pays de la Loire, France. **D.** Distance to the median in G2 Red Hawk rhizosphere in Michigan, USA. PERMDISP with Tukey HSD post-hoc test, * = p-value < 0.05, ** = p-value < 0.01. France n=5 plants per treatment, USA n=12 plants per treatment.

## Discussion

Our study brings insights into the complex responses of a legume-associated, below ground microbiome to drought exposure, and specifically tested for evidence of a of drought given consecutive exposure across two generations. While the droughts applied in our experiments negatively impacted the plants in both locations, notably, there was no strong evidence of a legacy effect of drought on the microbiome across two generations, with the notable exception of the Flavert rhizosphere microbiome. A feature of our experiment was the full-factorial two generation design with the use of two genotypes of common bean to understand the drought’s effects in two cultivars with different domestication histories (Beebe *et al*. 2001; Carrère *et al*. 2023). Flavert and Red Hawk genotypes originated from geographically isolated wild genotypes with distinct genetic pools, where Red Hawk traces its lineage to the Andean gene pool, while Flavert is a Flageolet bean accession from the European origin (Schmutz *et al*. 2014; Carrère *et al*. 2023). The literature has reported that the structure and composition of global bean rhizosphere microbiomes is influenced by the bean genotype (Pérez-Jaramillo *et al*. 2017, 2019). Thus, the distinct genetic lineages of the hosts likely drove differences in their associated microbiomes, which potentially explains the slightly different microbiome responses to drought observed between the two genotypes. It has been well-documented that host genetics can shape plant microbial structure and composition (Wagner *et al*. 2016; Brachi *et al*. 2022; Yue *et al*. 2023), that differences in the plant genome could affect the root exudation patterns and its rhizosphere microbiomes (Micallef, Shiaris and Colón-Carmona 2009; Sasse, Martinoia and Northen 2018; Wagner 2024). Thus, the contribution of plant host genotype in shaping plant-microbes interactions during drought stress is thought to be associated with root exudates (de Vries *et al*. 2020; Gu *et al*. 2024).

Over the two generations of drought exposure, and in both locations, the root and rhizosphere microbiomes of Red Hawk plants generally were resistant to drought. While water limitation can directly influence soil and rhizosphere microbiomes by shifting their structure and composition (Xu *et al*. 2018; Xie *et al*. 2021), there can also be indirect effects of drought on the microbiome as mediated by the host plant. Indirectly, drought can alter plant physiology and metabolism, subsequently affecting host-microbiome relationship and microbial responses to drought, with consequences for plant outcomes (Ait-El-Mokhtar, Meddich and Baslam 2023; Gholizadeh *et al*. 2024) and changes in rhizosphere microbial communities attributed to plant stress (Santos-Medellín *et al*. 2017; Veach *et al*. 2020). Even though the drought adaptation of Flavert bean has not been studied, our plant fitness data indicated a more strongly negative impact of drought on this genotype. There was an apparent effect of G1 drought on the rhizosphere microbiome of the Flavert plants in G2, suggesting a legacy effect of drought over generations, which was also reflected in the plant trait data. Given generational exposure to repeated droughts, plants may evolve and adapt by recruiting drought-specific microbes that are potentially beneficial for the host to withstand the stress (Terhorst, Lennon and Lau 2014; Naylor and Coleman-Derr 2018).

Red Hawk is a drought-susceptible red kidney bean genotype (Dramadri, Nkalubo and Kelly 2019), which is, in part, why it was chosen for this study. However, its relative sensitivity to drought was not reflected in the responses of its microbiomes, regardless of plant generation. This contrasts with other studies that have reported that the effect of drought on plant-associated microbiomes was more pronounced for drought-susceptible hosts (Bouasria *et al*. 2012; Martins *et al*. 2023). Here, we aimed to build upon our prior research to understand the responses of the common bean microbiome to drought exposure during the vegetative growth stage. Previously, we found no effect of drought in the microbiome of the seed endophytic compartment of Red Hawk but, instead, detected several stable bacterial taxa that were transmitted along parental lineages (Sulesky-Grieb *et al*. 2024), suggesting no yet-notable legacy effect of drought on the transmission of particular seed microbiome members. We also observed only a weak effect of short-term drought on the active rhizosphere microbiome structure of Red Hawk despite an obvious negative impact on the plants (e.g., wilting and final biomass reduction) (Bandopadhyay *et al*. 2024). Furthermore, while the host plant can be impacted by drought stress, the stress may not necessarily affect its associated microbiome (Babić *et al*. 2024). It has been reported that soil microbial communities have greater resistance and resilience to precipitation than plants (Cruz-Martínez *et al*. 2009; Curiel Yuste *et al*. 2014). A previous study has reported adaptation of rhizosphere microbiome under different magnitudes of drought (Dao *et al*.). Given that the plants responded negatively to the drought, the overall observation of resistance in the Red Hawk rhizosphere and root microbiomes is notable. More work is needed to determine whether the observed microbiome resistance to drought may help to offset detrimental effects on the host plant.

In addition, although other studies suggested that the effect of drought is stronger on endophytes compared to rhizosphere microbiomes due to immediate association with the host plant (Naylor *et al*. 2017; Santos-Medellín *et al*. 2017; Fitzpatrick *et al*. 2018), this did not apply to our study. There were no notable differences between rhizosphere and root-associated microbiomes (which included endophytes and rhizoplane together) in response to drought, with one exception: Flavert’s alpha diversity was decreased by drought in the rhizosphere of G2, but not in the root microbiomes.

We expected, and also detected, differences in soil microbiome composition from the different locations. Furthermore, we openly recognize that there were some minor differences in the execution of experiments across the two locations which are mostly due to differences in equipment between the two laboratories (e.g. growth chamber). These differences are an important feature of the work rather than a limitation. Parallel experiments are rarely executed and compared across different continents due to nuances in the experimental capacities available, but we suggest that these kinds of comparisons are needed to inform generalities and build theory towards prediction. Reporting the results from each location in separate manuscripts would miss the opportunity for comparison that our parallel experiments provide. As a contrast, in meta-analyses, generalities are deduced across different experimental designs, hosts, or production regions. While it is acknowledged that these “meta” comparisons are imperfect, it is also agreed that they provide insights by the merit of their bringing together.

Finally, it may be tempting to add more analyses, for example, to report differentially abundant ASVs and their taxonomies across the treatments. As the overarching trends show no notable differences across the treatments, and this could be perceived as hunting for a more exciting outcome. It could be that considering the active component of the microbiome, rather than the “total”, inclusive of DNA from active, inactive, and relic taxa, would have yielded a different perspective on the community response to drought (Bandopadhyay *et al*. 2024). We recommend future work to discriminate these components of the community, as the active taxa are those contributing to growth and plant-supportive functions.

To conclude, this study suggests that plant-associated microbiomes can remain stable (e.g., resistant) in response to stress despite negative impacts on the host, and that host genotype and soil origin can also mediate the microbiome responses. It did not provide evidence for any consistent or generalizable effect of drought legacy or multi-generational consequences of the drought on the below ground microbiome. Resistance of microbiomes is not often reported in the literature (potentially reflecting a reporting bias because it could be perceived as a “negative” result). However, resistance and resilience together contribute to the maintenance of stability in plant-microbiome systems, with consequences for agroecosystem responses to climate change. Thus, understanding the mechanisms that reinforce stability, either by resistance or resilience, and its implications for plant health is a critical next direction in plant-microbiome research.

## Acknowledgements

This work was supported by the United States Department of Agriculture (grant number 2019-67019-29305] to AS and MB, and by the Michigan State University Plant Resilience Institute to AS. This work was co-funded by the European Union [grant number ERC, MicroRescue, 101087042] to AS. Views and opinions expressed are however those of the author(s) only and do not necessarily reflect those of the European Union or the European Research Council. Neither the European Union nor the granting authority can be held responsible for them. AS acknowledges support from the United States Department of Agriculture National Institute of Food and Agriculture and Michigan State University AgBioResearch, and the Centre National de la Recherche Scientifique (CNRS), France. The Angers Plant Phenotyping Platform PHENOTIC (DOI: 10.17180/ykbz-2v85) is acknowledged for the production and phenotyping of plants.

