## Supplemental Information for "Evaluating the legacy of drought exposure on root and rhizosphere bacterial microbiomes over two plant generations"

Supplemental Material

### Table S1. Parameters used in 16S rRNA amplicon sequence processing

| Dataset | Dada2 trim parameters | Taxonomy assignment | Decontam parameters | Rarefaction |
|---|---|---|---|---|
| **Michigan, USA Roots 16S** | --p-trunc-len-f 114 \ --p-trunc-len-r 161 \ | silva-138-99-515-806-nb-classifier.qza | Threshold 0.1 | 20,976 |
| **Michigan, USA Rhizosphere 16S** | --p-trunc-len-f 103 \ --p-trunc-len-r 172 \ | silva-138-99-515-806-nb-classifier.qza | Threshold 0.1 | 20,976 |
| **Pays de la Loire, France Root and Rhizosphere 16S - Generation 1** | --p-trunc-len-f 200 \ --p-trunc-len-r 180- \ | Silva 132 99% | Threshold 0.2 | 2,500 |
| **Pays de la Loire, France Root and Rhizosphere 16S - Generation 2** | --p-trunc-len-f 200 \ --p-trunc-len-r 130- \ | Silva 132 99% | Threshold 0.2 | 2,500 |

**Figure S1.** Rarefaction curves of number of amplicon sequence variants (ASVs) "Species Richness" by number of reads ("Sample Size") for each microbiome profile generated by 16S rRNA amplicon sequencing. Rhizosphere samples are provided in pink and root in blue. MSU shows samples from Michigan State University, Michigan, USA, and INRAE shows samples from Pays de la Loire, France. Vertical lines show the subsampling depth applied for alpha and beta diversity analyses (20,976 reads per sample for Michigan, and 2,500 reads per sample for Pays de la Loire). Note differences in both y- and x-axis ranges across the two panels.

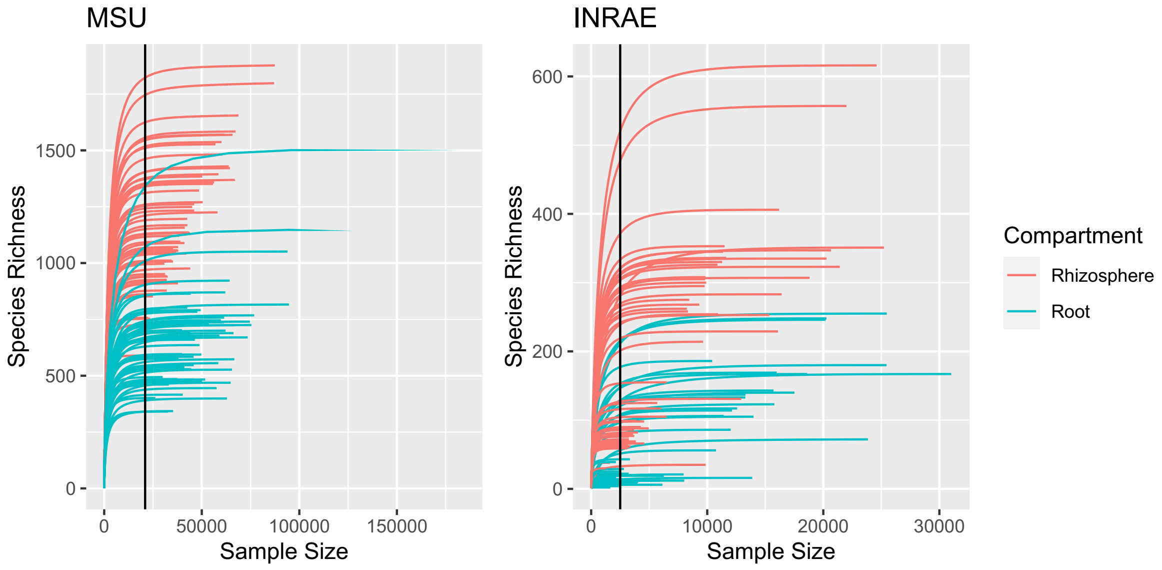

**Table S2. Effects of drought treatment, plant genotype, and their interaction on generation 1 (G1) plant phenotypes (pod count, seed count, above-ground and root dry mass, photosynthetic rate, and stomatal conductance) were assessed by two-way ANOVA (Pays de la Loire, France) and Welch's two-sample t-test (Michigan, USA) (df = degrees of freedom, F = F-statistic, t = t-statistic, P = P-value, values P<0.05 are indicated in bold)**

| Two-way ANOVA | Pays de la Loire (France) | | | Welch's Two Sample t-test | Michigan (USA) | | |
|---|---|---|---|---|---|---|---|
| | df | F | P | | df | t | P |
| **Pod count** | | | | **Pod count** | | | |
| *Genotype* | 1 | 355.618 | **2.36e-12** | *G1 drought* | 5.96 | 0 | 1 |
| *G1 drought* | 1 | 58.066 | **1.04e-06** | | | | |
| *Genotype x G1 drought* | 1 | 0.776 | 0.391 | | | | |
| **Seed count** | | | | **Seed count** | | | |
| *Genotype* | 1 | 100.292 | **2.69e-08** | *G1 drought* | 7.6 | 0.071 | 0.946 |
| *G1 drought* | 1 | 0.082 | 0.778 | | | | |
| *Genotype x G1 drought* | 1 | 0.002 | 0.968 | | | | |
| **Above-ground mass** | | | | **Above-ground mass** | | | |
| *Genotype* | 1 | 3.304 | 0.0891 | *G1 drought* | 7.96 | 2.497 | **0.037** |
| *G1 drought* | 1 | 5.058 | **0.04** | | | | |
| *Genotype x G1 drought* | 1 | 0.459 | 0.5084 | | | | |
| **Root mass** | | | | **Root mass** | | | |
| *Genotype* | 1 | 7.648 | **0.0138** | *G1 drought* | 6.37 | 1.189 | 0.277 |
| *G1 drought* | 1 | 2.377 | 0.1426 | | | | |
| *Genotype x G1 drought* | 1 | 0.017 | 0.8985 | | | | |
| | | | | **Photosynthetic rate** | | | |
| | | | | *G1 drought* | 5.71 | 3.283 | **0.018** |
| | | | | **Stomatal conductance** | | | |
| | | | | *G1 drought* | 6.16 | 2.816 | **0.029** |

**Table S3. Effects of drought treatment on generation 1 (G1) plant phenotypes (pod count, seed count, above-ground and root dry mass) within each genotype in Pays de la Loire, France were assessed by Welch's two-sample t-test (df = degrees of freedom, t = t-statistic, P = P-value, values P<0.05 are indicated in bold)**

| Welch's two-sample t-test | Pays de la Loire (France) | | | | | |
|---|---|---|---|---|---|---|
| | Flavert | | | Red Hawk | | |
| | df | t | P | df | t | P |
| **Pod count** | | | | | | |
| *G1 drought* | 7.5955 | 7.0602 | **0.0001363** | 6.6301 | 4.264 | **0.00422** |
| **Seed count** | | | | | | |
| *G1 drought* | 7.6164 | 0.13001 | 0.8999 | 7.2645 | 0.49237 | 0.637 |
| **Above-ground mass** | | | | | | |
| *G1 drought* | 4.1567 | 1.7858 | 0.146 | 4.4113 | 1.4781 | 0.207 |
| **Root mass** | | | | | | |
| *G1 drought* | 5.8955 | 0.94479 | 0.3819 | 7.9852 | 1.5139 | 0.1686 |

**Table S4. Effects of drought treatment (G1 drought = parent drought treatment, G2 drought = current drought treatment), plant genotype, and their interactions on generation 2 (G2) plant phenotypes (pod count, seed count, shoot and root dry mass, photosynthetic rate, and stomatal conductance) were assessed by three-way ANOVA (Pays de la Loire, France) and two-way ANOVA (Michigan, US) (df = degrees of freedom, F = F-statistic, P = P-value, values P<0.05 are indicated in bold)**

| **Three-way ANOVA** | **Pays de la Loire (France)** | | | **Two-way ANOVA** | **Michigan (USA)** | | |
|---|---|---|---|---|---|---|---|
| | **df** | **F** | **P** | | **df** | **F** | **P** |
| **Pod count** | | | | **Pod count** | | | |
| *Genotype* | 1 | 419.767 | **< 2e-16** | *G1 drought* | 1 | 2.038 | 0.161 |
| *G1 drought* | 1 | 0.419 | 0.522248 | *G2 drought* | 1 | 0.233 | 0.632 |
| *G2 drought* | 1 | 111.674 | **5.77e-12** | *G1 drought x G2 drought* | 1 | 0.442 | 0.510 |
| *Genotype x G1 drought* | 1 | 0.419 | 0.522248 | | | | |
| *Genotype x G2 drought* | 1 | 16.791 | **0.000266** | | | | |
| *G1 drought x G2 drought* | 1 | 0.047 | 0.830617 | | | | |
| *Genotype x G1 drought x G2 drought* | 1 | 1.163 | 0.288949 | | | | |
| **Seed count** | | | | **Seed count** | | | |
| *Genotype* | 1 | 921.891 | **< 2e-16** | *G1 drought* | 1 | 0.013 | 0.909 |
| *G1 drought* | 1 | 2.189 | 0.148815 | *G2 drought* | 1 | 0.791 | 0.379 |
| *G2 drought* | 1 | 169.851 | **2.42e-14** | *G1 drought x G2 drought* | 1 | 1.362 | 0.250 |
| *Genotype x G1 drought* | 1 | 0.045 | 0.833961 | | | | |
| *Genotype x G2 drought* | 1 | 17.275 | **0.000225** | | | | |
| *G1 drought x G2 drought* | 1 | 0.124 | 0.726971 | | | | |
| *Genotype x G1 drought x G2 drought* | 1 | 2.625 | 0.114986 | | | | |
| **Above-ground mass** | | | | **Above-ground mass** | | | |
| *Genotype* | 1 | 3.061 | 0.0898 | *G1 drought* | 1 | 0.003 | 0.958 |
| *G1 drought* | 1 | 8.953 | **0.0053** | *G2 drought* | 1 | 0.180 | 0.674 |
| *G2 drought* | 1 | 368.450 | **<2e-16** | *G1 drought x G2 drought* | 1 | 1.221 | 0.275 |
| *Genotype x G1 drought* | 1 | 3.870 | 0.0579 | | | | |
| *Genotype x G2 drought* | 1 | 1.369 | 0.2507 | | | | |
| *G1 drought x G2 drought* | 1 | 0.001 | 0.9754 | | | | |
| *Genotype x G1 drought x G2 drought* | 1 | 0.227 | 0.6371 | | | | |
| **Root mass** | | | | **Root mass** | | | |
| *Genotype* | 1 | 8.192 | **0.007361** | *G1 drought* | 1 | 0.390 | 0.536 |
| *G1 drought* | 1 | 3.585 | 0.067364 | *G2 drought* | 1 | 0.070 | 0.793 |
| *G2 drought* | 1 | 14.801 | **0.000537** | *G1 drought x G2 drought* | 1 | 0.027 | 0.869 |
| *Genotype x G1 drought* | 1 | 2.416 | 0.129973 | | | | |
| *Genotype x G2 drought* | 1 | 13.645 | **0.000820** | | | | |
| *G1 drought x G2 drought* | 1 | 0.001 | 0.974010 | | | | |
| *Genotype x G1 drought x G2 drought* | 1 | 0.692 | 0.411673 | | | | |
| | | | | **Photosynthetic rate** | | | |

| | | | |
|---|---|---|---|
| *G1 drought* | 1 | 0.009 | 0.926087 |
| *G2 drought* | 1 | 18.425 | **0.000642** |
| *G1 drought x G2 drought* | 1 | 0.211 | 0.652636 |
| **Stomatal Conductance** | | | |
| *G1 drought* | 1 | 0.177 | 0.68 |
| *G2 drought* | 1 | 46.803 | **5.59e-06** |
| *G1 drought x G2 drought* | 1 | 0.154 | 0.70 |

**Table S5. Effects of G1 drought (parent drought treatment), G2 drought (current drought treatment), and their interaction on generation 2 (G2) plant phenotypes (pod count, seed count, above-ground and root dry mass) within each genotype in Pays de la Loire, France were assessed by two-way ANOVA (df = degrees of freedom, F = F-statistic, P = P-value, values P<0.05 are indicated in bold)**

| Two-way ANOVA | Pays de la Loire (France) | | | | | |
|---|---|---|---|---|---|---|
| | Flavert | | | Red Hawk | | |
| | df | F | P | df | F | P |
| **Pod count** | | | | | | |
| *G1 drought* | 1 | 0 | 1 | 1 | 1.125 | 0.305 |
| *G2 drought* | 1 | 85.630 | **7.99e-08** | 1 | 28.125 | **7.14e-05** |
| *G1 drought x G2 drought* | 1 | 0.296 | 0.594 | 1 | 1.125 | 0.305 |
| **Seed count** | | | | | | |
| *G1 drought* | 1 | 0.83 | 0.376 | 1 | 2.893 | 0.108 |
| *G2 drought* | 1 | 85.79 | **7.89e-08** | 1 | 141.750 | **2.31e-09** |
| *G1 drought x G2 drought* | 1 | 1.13 | 0.304 | 1 | 2.893 | 0.108 |
| **Above-ground mass** | | | | | | |
| *G1 drought* | 1 | 0.410 | 0.531 | 1 | 17.106 | **0.000776** |
| *G2 drought* | 1 | 161.863 | **8.78e-10** | 1 | 225.981 | **7.4e-11** |
| *G1 drought x G2 drought* | 1 | 0.077 | 0.784 | 1 | 0.179 | 0.677861 |
| **Root mass** | | | | | | |
| *G1 drought* | 1 | 8.434 | **0.0104** | 1 | 0.044 | 0.835701 |
| *G2 drought* | 1 | 0.017 | 0.8989 | 1 | 21.952 | 0.000248 |
| *G1 drought x G2 drought* | 1 | 0.453 | 0.5105 | 1 | 0.289 | 0.598514 |

**Table S6. Effects of drought treatment, plant genotype, and their interaction on root and rhizosphere alpha diversity (richness) of generation 1 (G1) plant were assessed by two-way ANOVA (Pays de la Loire, France) and Welch's two-sample t-test (Michigan, USA) (df = degrees of freedom, F = F-statistic, t = t-statistic, P = P-value, values P<0.05 are indicated in bold)**

| **Two-way ANOVA** | **Pays de la Loire (France)** | | | **Welch's Two Sample t-test** | **Michigan (USA)** | | |
|---|---|---|---|---|---|---|---|
| | **df** | **F** | **P** | | **df** | **t** | **P** |
| **Root** | | | | **Root** | | | |
| *Genotype* | 1 | 7.651 | **0.0138** | *G1 drought* | 6.87 | 0.055 | 0.958 |
| *G1 drought* | 1 | 3.303 | 0.0879 | | | | |
| *Genotype x G1 drought* | 1 | 3.534 | 0.0785 | | | | |
| **Rhizosphere** | | | | **Rhizosphere** | | | |
| *Genotype* | 1 | 3.395 | 0.084 | *G1 drought* | 4.067 | 0.792 | 0.472 |
| *G1 drought* | 1 | 0.271 | 0.61 | | | | |
| *Genotype x G1 drought* | 1 | 3.267 | 0.0895 | | | | |

**Table S7. Effects of drought treatment (G1 drought = parent drought treatment, G2 drought = current drought treatment), plant genotype, and their interactions on root and rhizosphere alpha diversity (richness) of generation 2 (G2) plant were assessed by three-way ANOVA (Pays de la Loire, France) and two-way ANOVA (Michigan, USA) (df = degrees of freedom, F = F-statistic, P = P-value, values P<0.05 are indicated in bold)**

| Three-way ANOVA | Pays de la Loire (France) | | | Two-way ANOVA | Michigan (USA) | | |
|---|---|---|---|---|---|---|---|
| | df | F | P | | df | F | P |
| **Root** | | | | **Root** | | | |
| *Genotype* | 1 | 2.490 | 0.1267 | *G1 drought* | 1 | 0.151 | 0.699 |
| *G1 drought* | 1 | 0.228 | 0.6372 | *G2 drought* | 1 | 2.422 | 0.127 |
| *G2 drought* | 1 | 7.023 | **0.0135** | *G1 drought x G2 drought* | 1 | 1.452 | 0.235 |
| *Genotype x G1 drought* | 1 | 0.650 | 0.4275 | | | | |
| *Genotype x G2 drought* | 1 | 1.556 | 0.2234 | | | | |
| *G1 drought x G2 drought* | 1 | 1.188 | 0.2857 | | | | |
| *Genotype x G1 drought x G2 drought* | 1 | 0.034 | 0.8557 | | | | |
| **Rhizosphere** | | | | **Rhizosphere** | | | |
| *Genotype* | 1 | 4.983 | **0.03378** | *G1 drought* | 1 | 0.685 | 0.412 |
| *G1 drought* | 1 | 9.132 | **0.00532** | *G2 drought* | 1 | 0.729 | 0.397 |
| *G2 drought* | 1 | 0.266 | 0.61032 | *G1 drought x G2 drought* | 1 | 4.145 | **0.047** |
| *Genotype x G1 drought* | 1 | 1.336 | 0.25748 | | | | |
| *Genotype x G2 drought* | 1 | 0.574 | 0.45490 | | | | |
| *G1 drought x G2 drought* | 1 | 1.136 | 0.29555 | | | | |
| *Genotype x G1 drought x G2 drought* | 1 | 2.506 | 0.12463 | | | | |

**Table S8. Effects of G1 drought (parent drought treatment), G2 drought (current drought treatment), and their interaction on generation 2 (G2) on root and rhizosphere alpha diversity (richness) of generation 2 (G2) plant within each genotype in Pays de la Loire, France were assessed by two-way ANOVA (df = degrees of freedom, F = F-statistic, P = P-value, values P<0.05 are indicated in bold)**

| Two-way ANOVA | Pays de la Loire (France) | | | | | |
|---|---|---|---|---|---|---|
| | Flavert | | | Red Hawk | | |
| | **df** | **F** | **P** | **df** | **F** | **P** |
| **Root** | | | | | | |
| *G1 drought* | 1 | 1.470 | 0.245 | 1 | 0.001 | 0.976 |
| *G2 drought* | 1 | 0.456 | 0.511 | 1 | 8.734 | **0.012** |
| *G1 drought x G2 drought* | 1 | 0.300 | 0.593 | 1 | 1.158 | 0.303 |
| **Rhizosphere** | | | | | | |
| *G1 drought* | 1 | 25.631 | **0.000173** | 1 | 1.148 | 0.302 |
| *G2 drought* | 1 | 1.651 | 0.219691 | 1 | 0.016 | 0.901 |
| *G1 drought x G2 drought* | 1 | 2.583 | 0.130349 | 1 | 1.217 | 0.289 |

**Table S9. Effects of drought treatment, plant genotype, and their interaction on beta diversity (Bray-Curtis distance matrix) of generation 1 (G1) plant were assessed by PERMANOVA (df = degrees of freedom, $R^2$ = coefficient of determination, F = F-statistic, P = P-value, number of permutations = 9999, values P<0.05 are indicated in bold)**

| PERMANOVA | Pays de la Loire (France) | | | | PERMANOVA | Michigan (USA) | | | |
|---|---|---|---|---|---|---|---|---|---|
| | df | $R^2$ | F | P | | df | $R^2$ | F | P |
| **Root** | | | | | **Root** | | | | |
| *Genotype* | 1 | 0.48604 | 17.0862 | **0.0001** | *G1 drought* | 1 | 0.15159 | 1.4294 | 0.1398 |
| *G1 drought* | 1 | 0.03054 | 1.0735 | 0.3092 | | | | | |
| *Genotype x G1 drought* | 1 | 0.02829 | 0.9944 | 0.3329 | | | | | |
| **Rhizosphere** | | | | | **Rhizosphere** | | | | |
| *Genotype* | 1 | 0.06119 | 1.2232 | 0.2185 | *G1 drought* | 1 | 0.10282 | 0.9168 | 0.4316 |
| *G1 drought* | 1 | 0.08592 | 1.7175 | **0.0241** | | | | | |
| *Genotype x G1 drought* | 1 | 0.05246 | 1.0487 | 0.2943 | | | | | |

**Table S10. Effects of drought treatment and their interaction on beta diversity (Bray-Curtis distance matrix) of generation 1 (G1) plant within each genotype in Pays de la Loire, France were assessed by PERMANOVA (df = degrees of freedom, $R^2$ = coefficient of determination, F = F-statistic, P = P-value, number of permutations = 9999, values P<0.05 are indicated in bold)**

| PERMANOVA | Pays de la Loire (France) | | | | | | | |
|---|---|---|---|---|---|---|---|---|
| | Flavert | | | | Red Hawk | | | |
| | df | $R^2$ | F | P | df | $R^2$ | F | P |
| **Root** | | | | | | | | |
| *G1 drought* | 1 | 0.11572 | 1.0469 | 0.3451 | 1 | 0.11197 | 1.0087 | 0.3858 |
| **Rhizosphere** | | | | | | | | |
| *G1 drought* | 1 | 0.16482 | 1.5787 | **0.021** | 1 | 0.13206 | 1.2172 | 0.1011 |

**Table S11. Effects of drought treatment (G1 drought = parent drought treatment, G2 drought = current drought treatment), plant genotype, and their interactions on beta diversity (Bray-Curtis distance matrix) of generation 2 (G2) plant were assessed by PERMANOVA (df = degrees of freedom, $R^2$ = coefficient of determination, F = F-statistic, P = P-value, number of permutations = 9999, values P<0.05 are indicated in bold).**

| PERMANOVA | Pays de la Loire (France) | | | | PERMANOVA | Michigan (USA) | | | |
|---|---|---|---|---|---|---|---|---|---|
| | df | $R^2$ | F | P | | df | $R^2$ | F | P |
| **Root** | | | | | **Root** | | | | |
| *Genotype* | 1 | 0.30600 | 15.4685 | **0.0006** | *G1 drought* | 1 | 0.01702 | 0.7980 | 0.2832 |
| *G1 drought* | 1 | 0.01027 | 0.5190 | 0.5053 | *G2 drought* | 1 | 0.02670 | 1.2517 | **0.0290** |
| *G2 drought* | 1 | 0.00607 | 0.3067 | 0.6443 | *G1 drought x G2 drought* | 1 | 0.01767 | 0.8283 | 0.2239 |
| *Genotype x G1 drought* | 1 | 0.01047 | 0.5292 | 0.4970 | | | | | |
| *Genotype x G2 drought* | 1 | 0.02327 | 1.1762 | 0.2767 | | | | | |
| *G1 drought x G2 drought* | 1 | 0.12623 | 6.3810 | **0.0111** | | | | | |
| *Genotype x G1 drought x G2 drought* | 1 | 0.00337 | 0.1704 | 0.8049 | | | | | |
| **Rhizosphere** | | | | | **Rhizosphere** | | | | |
| *Genotype* | 1 | 0.06125 | 2.5661 | **0.0007** | *G1 drought* | 1 | 0.01450 | 0.6848 | 0.5682 |
| *G1 drought* | 1 | 0.06888 | 2.8856 | **0.0002** | *G2 drought* | 1 | 0.01542 | 0.7281 | 0.3975 |
| *G2 drought* | 1 | 0.05483 | 2.2972 | **0.0031** | *G1 drought x G2 drought* | 1 | 0.01708 | 0.8065 | 0.1801 |
| *Genotype x G1 drought* | 1 | 0.06560 | 2.7483 | **0.0006** | | | | | |
| *Genotype x G2 drought* | 1 | 0.01959 | 0.8206 | 0.6887 | | | | | |
| *G1 drought x G2 drought* | 1 | 0.02926 | 1.2259 | 0.2074 | | | | | |
| *Genotype x G1 drought x G2 drought* | 1 | 0.03224 | 1.3505 | 0.1358 | | | | | |

**Table S12. Effects of G1 drought (parent drought treatment), G2 drought (current drought treatment), and their interactions on beta diversity (Bray-Curtis distance matrix) of generation 2 (G2) plant within each genotype in Pays de la Loire, France were assessed by PERMANOVA (df = degrees of freedom, $R^2$ = coefficient of determination, F = F-statistic, P = P-value, number of permutations = 9999, values P<0.05 are indicated in bold).**

| PERMANOVA | Pays de la Loire (France) | | | | | | | |
|---|---|---|---|---|---|---|---|---|
| | Flavert | | | | Red Hawk | | | |
| | df | $R^2$ | F | P | df | $R^2$ | F | P |
| **Root** | | | | | | | | |
| *G1 drought* | 1 | 0.03099 | 0.6254 | 0.4759 | 1 | 0.02719 | 0.4127 | 0.5974 |
| *G2 drought* | 1 | 0.03024 | 0.6102 | 0.4803 | 1 | 0.05641 | 0.8560 | 0.3836 |
| *G1 drought x G2 drought* | 1 | 0.24490 | 4.9412 | **0.0272** | 1 | 0.12569 | 1.9075 | 0.1750 |
| **Rhizosphere** | | | | | | | | |
| *G1 drought* | 1 | 0.20094 | 4.3202 | **0.0001** | 1 | 0.05668 | 0.9956 | 0.4454 |
| *G2 drought* | 1 | 0.07582 | 1.6302 | **0.0497** | 1 | 0.08992 | 1.5795 | 0.0525 |
| *G1 drought x G2 drought* | 1 | 0.07205 | 1.5491 | 0.0536 | 1 | 0.05635 | 0.9898 | 0.4634 |

**Table S13. Beta dispersion analysis of generation 1 (G1) plants were assessed by PERMDISP (df = degrees of freedom, F = F-statistic, P = P-value, number of permutations = 999, values P<0.05 are indicated in bold)**

| PERMADISP | Pays de la Loire (France) | | | | | | PERMADISP | Michigan (USA) | | |
|---|---|---|---|---|---|---|---|---|---|---|
| | Flavert | | | Red Hawk | | | | Red Hawk | | |
| | df | F | P | df | F | P | | df | F | P |
| **Root** | | | | | | | | | | |
| *G1 drought* | 1 | 0.798 | 0.448 | 1 | 0.6708 | 0.494 | *G1 drought* | 1 | 0.5402 | 0.488 |
| **Rhizosphere** | | | | | | | | | | |
| *G1 drought* | 1 | 4.0062 | **0.046** | 1 | 1.3403 | 0.301 | *G1 drought* | 1 | 0.0581 | 0.817 |

**Table S14. Beta dispersion analysis of generation 2 (G2) plants were assessed by PERMDISP (df = degrees of freedom, F = F-statistic, P = P-value, number of permutations = 999, values P<0.05 are indicated in bold)**

| PERMADISP | Pays de la Loire (France) | | | | | | PERMADISP | Michigan (USA) | | |
|---|---|---|---|---|---|---|---|---|---|---|
| | Flavert | | | Red Hawk | | | | Red Hawk | | |
| | df | F | P | df | F | P | | df | F | P |
| **Root** | | | | | | | **Root** | | | |
| *G1 drought* | 1 | 0.0886 | 0.759 | 1 | 0.5329 | 0.478 | *G1 drought* | 1 | 1.1104 | 0.292 |
| *G2 drought* | 1 | 0.0091 | 0.941 | 1 | 1.8366 | 0.198 | *G2 drought* | 1 | 1.0537 | 0.309 |
| *G1 X G2 drought* | 3 | 0.8629 | 0.496 | 3 | 1.6994 | 0.211 | *G1 X G2 drought* | 3 | 1.2683 | 0.265 |
| **Rhizosphere** | | | | | | | **Rhizosphere** | | | |
| *G1 drought* | 1 | 0.5645 | 0.488 | 1 | 0.0022 | 0.96 | *G1 drought* | 1 | 2.6643 | 0.113 |
| *G2 drought* | 1 | 4.0549 | **0.048** | 1 | 1.8265 | 0.181 | *G2 drought* | 1 | 5.1359 | **0.028** |
| *G1 X G2 drought* | 3 | 4.1156 | **0.02** | 3 | 0.2518 | 0.843 | *G1 X G2 drought* | 3 | 1.6794 | 0.179 |
